# A map of climate change-driven natural selection in *Arabidopsis thaliana*

**DOI:** 10.1101/321133

**Authors:** Moises Exposito-Alonso, 500 Genomes Field Experiment Team, Hernán A. Burbano, Oliver Bossdorf, Rasmus Nielsen, Detlef Weigel

**Author notes:** Current address: Department of Integrative Biology, University of California Berkeley, Berkeley, CA 94720, USA.

## Abstract

Through the lens of evolution, climate change is an agent of natural selection that forces populations to change and adapt, or face extinction. Current assessments of the risk of biodiversity associated with climate change^1^, however, do not typically take into account the genetic makeup of populations and how natural selection impacts it^2^. We made use of the extensive genome information in *Arabidopsis thaliana* and measured how rainfall-manipulation affected the fitness of 517 natural lines grown in Spain and Germany. This allowed us to directly infer selection along the genome^3^. Natural selection was particularly strong in the hot-dry Spanish location, killing 63% of lines and significantly changing the frequency of ~5% of all genome-wide variants. A significant portion of this climate-driven natural selection over variants was predictable from signatures of local adaptation (R^2^=29-52%), as genetic variants found in geographic areas with climates more similar to the experimental sites were positively selected. Field-validated predictions across the species range indicated that Mediterranean and Western Siberian populations — at the edges of the species’ environmental limits — currently experience the strongest climate-driven selection. With more frequent droughts and rising temperatures in Europe^4^, we forecast an increase in directional natural selection moving northwards from the southern end, and putting many native *A. thaliana* populations at evolutionary risk.

---

To predict the future impact of climate change on biodiversity, the typical starting point has been to study the limits of climate tolerance inferred from a species’ present geographic distributions. These tolerances are usually treated as static over time, and risks are assessed based on whether the geographic areas with climates within the tolerance limits will shrink^1^ or shift faster than the species can migrate^1,5^. However, these approaches do not account for within-species genetic variation, nor for how natural selection causes species to genetically change and adapt over time^2^.

Thanks to species-wide genome selection scans^6,7^ as well as genome associations with climate of origin^8–12^, we increasingly understand the genomic basis of past selection and climate adaptation. The best way to quantify current selection in a specific environment is provided by field experiments in which multiple genotypes of a species are grown together^13,14^ and relative fitness is directly associated with genetic variation^3,15^. We combined such knowledge with global climate change projections to predict the “evolutionary impact” of climate change on a species, i.e. how much local populations across the species range will need to genetically change and re-adapt to future climates.

To study natural selection in the annual plant *A. thaliana*, we performed two common garden experiments for one generation in two climatically distinct field stations, at the warm edge of the species distribution in Madrid (Spain, <u>40.40805ºN -3.83535ºE</u>), and closer to the distribution center in Tübingen (Germany, <u>48.545809ºN 9.042449ºE</u>) (Fig. 1, for details see Supplemental Appendix II). At each site, we simulated high precipitation typical of Germany, and low precipitation typical of Spain (we used four flooding tables with a split replicated design of two wet and two dry treatments, each with four spatial blocks, identically replicated at both sites, see Fig. SII.2, Table SII.1). In fall of 2015 we sowed over 300,000 seeds of 517 randomized natural lines capturing species-wide genomic diversity^16^ (<u>Dataset 1-2</u>). For each line, we prepared seven pots in which only a single plant was retained after germination *in situ*, and five pots with exactly 30 seeds that were allowed to germinate and grow without intervention throughout the experiment. At the end of the experiment in June 2016, we had collected data from 23,154 pots, consisting of survival to the reproductive stage and the number of seeds per surviving plant (fecundity), and their product, lifetime fitness (<u>Dataset 3-4</u>). Heritability of fitness varied across environments and between survival and fecundity. It was generally highest in the most stressful environment (H^2^_survival_=0.551; Table SI.3), i.e. in Spain under low precipitation and at high plant density. In this environment, only 193 of the 517 accessions survived, whereas in Germany at least a few plants of each accession reproduced (Table SI.1).

**Fig. 1.**
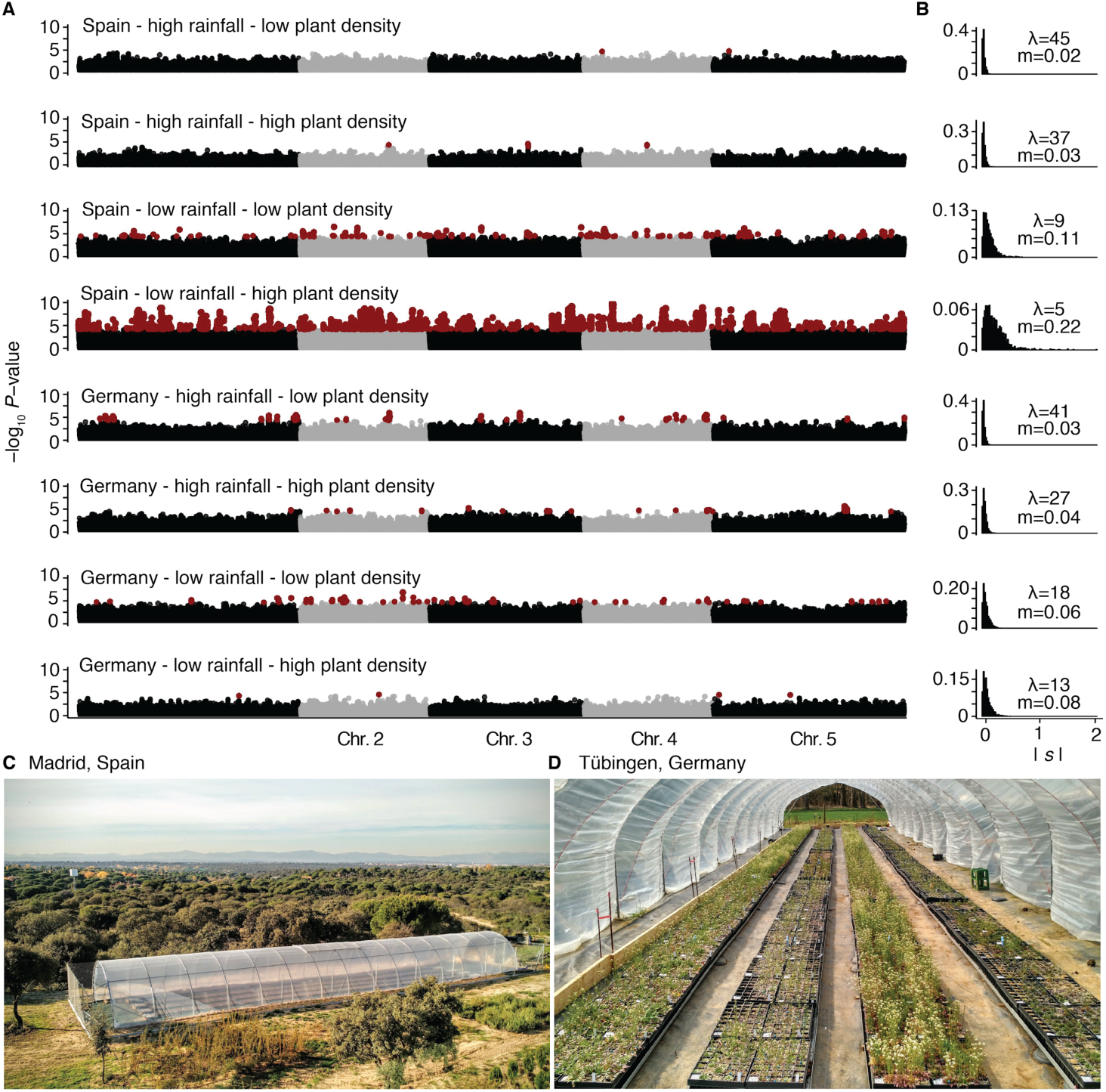
A genome map of total selection coefficients. (A) Manhattan plots of SNPs significantly associated with relative lifetime fitness in eight different environments. SNPs significant after FDR (black and grey) or Bonferroni correction (red) are shown. For genome-wide scans of survival and fecundity see Fig. SI.4 and SI.5. (B) Distribution of absolute total selection coefficients |*S*| per experiment. λ denotes maximum likelihood-inferred parameter of an exponential distribution, and *m* denotes the mean total selection coefficient. (C) Aerial picture of the experimental site in Spain. (D) Close-up picture inside the opened foil tunnel in Germany.

In each experimental environment, we quantified natural selection over minor alleles, what we call *total selection coefficient* (*s*, ref. ^17^), based on the difference in relative fitness of lines with the minor and the major allele at each genomic position (1,353,386 biallelic SNPs in 515 lines using a Genome-Wide Association [GWA] approach with Linear Models [LM-GEMMA] ref. ^18^, see Supplemental Appendix I section IV). In real populations, especially in selfing species with extensive population structure, alleles linked to causal variants cannot escape the consequences of selection, that is, an increase or decrease in frequency or even fixation or loss - a phenomenon behind background selection and genetic hitchhiking^19–21^. With *s* we capture the realized selection affecting each SNP’s minor allele, which results from the combination of selection acting directly on the focal allele, and the indirect effects due to selection on causal SNPs that are in LD with the focal variant. This approach best reflects an increase or decrease in frequency of each SNP in the population after one generation of selection (see simulations Fig. SI.15). The inferred *s* coefficient was significantly different from zero in at least one of the eight environments for 6,538 SNPs, under a Bonferroni corrected threshold (*P*<7×10^−7^), or 421,962 SNPs, under a 0.05 threshold after Benjamini & Hochberg FDR correction (Fig. 1, Table SI.3). If we considered our experiment as a naturally evolving population, the top 1*% s* coefficients in Spain would cause allele frequencies to change in the range of 12 to 24%, and another ~1% of all genome-wide alleles (n=12,179) would become fixed in one generation (this is due to many accessions did not survive whatsoever). In the benign high-precipitation environment of Germany, the top 1% *s* coefficients would push allele frequencies to change by 1.2 to 2.6%, and no allele would become fixed (Table SI.4, Fig. SI.9, Supplemental Appendix I section IV).

We were mainly interested in how a population’s genetic makeup changes in response to natural selection, but we were also curious what fraction of the alleles that change in frequency might be under indirect versus direct selection. To this end, we carried out another genome-wide association removing indirect LD-driven effects^17,18^, commonly known as population structure correction, using Bayesian Sparse Linear Mixed Model (BSLMM-GEMMA, ref. ^18^, see Supplemental Appendix I section IV). This analysis estimated that the most likely number of causal loci (*nγ*) was in the range of 7 to 89 per experiment (Table SI.3). This confirmed our expectation that the vast majority of detected alleles (Fig. 1) experienced indirect natural selection^17,20^. In agreement with BSLMM analyses being designed to find SNPs with potential direct effects on fitness, the inferred causal alleles were also more likely to be nonsynonymous rather than synonymous mutations (N=368 alleles with inclusion probability *γ*>5% in any of the experiments, Fisher’s Exact test Odds Ratio [OR] =1.32, *P*=1.18×10^−5^).

An important question is whether alleles that are beneficial in one environment are beneficial, neutral or even detrimental in another environment. We found that alleles positively selected under low precipitation tended to be negatively selected under high precipitation, and vice versa, so-called antagonistic pleiotropy^22^ (Fig. 2, Fisher’s exact test OR >1.31, *P*<4×10^−24^) — an observation that is particularly clear when comparing the two most “natural” conditions, low precipitation in Spain and high precipitation in Germany (OR=6.72). In contrast, when we compared the same precipitation condition between the two locations, selection was either in the same direction (0.23<Pearson’s r<0.51), or there was selection in one environment and neutrality in the other, so-called conditional neutrality (All OR<1, *P*<10^−16^).

**Fig. 2.**
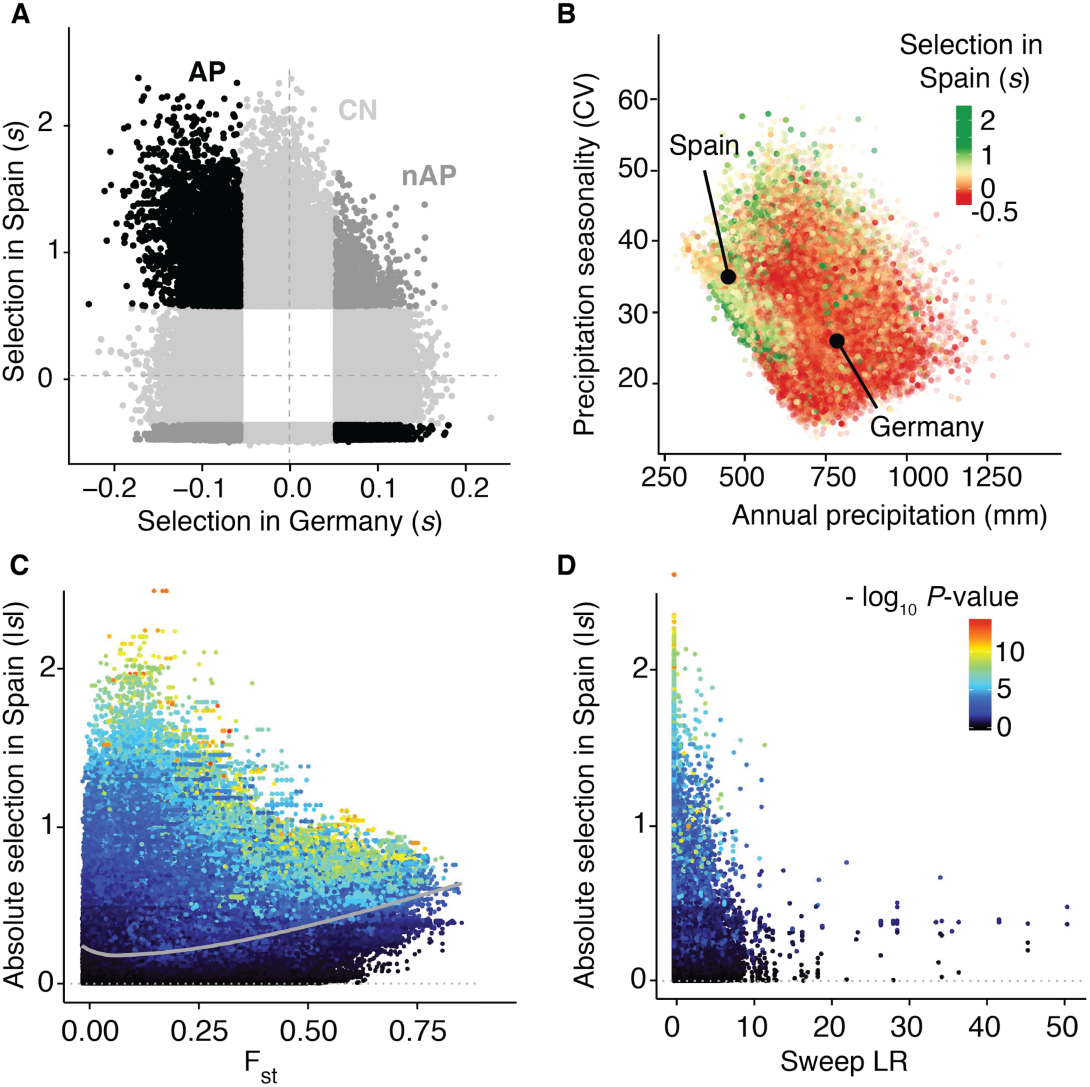
Selection trade-offs and the signal of environmental local adaptation. (A) 5% extreme tails of total selection coefficients across two contrasting environments; Spain with low precipitation and high population density, and Germany with high precipitation and low population density. In light grey are conditionally neutral alleles for either environment (CN, n=265,436), in black are alleles behaving as antagonistic pleiotropic (AP, n=20,503), and in dark grey are alleles behaving as non-antagonistic pleiotropic (nAP, n=9,681). (B) Mean annual precipitation and precipitation seasonality in the geographic areas of origin of SNPs (n=1,353,386 SNPs). Black circles indicate the average climate values at our Spanish (left) and German experimental sites (right). (C) Relationship between field absolute total selection coefficients and *F_ST_* values across 11 lineages, and (D) the likelihood ratio of selective sweeps (n=1,353,386 SNPs).

If the above patterns mimic natural selection driven by climate in the wild, we should be able to find aligned footprints of past selection and adaptation along the genome of our populations. To this end, we searched for selective sweeps^6^, outlier allele frequency differentiation between eleven *A. thaliana* genetic groups^8^ (*F_ST_*), and climate-allele associations^10,11^ (GWA with 1960-1990 average climate of origin, worldclim.org, ref. ^23^) (see Supplemental Appendix I section II, V and VI). Bonferroni-significant *s* had a higher average *F_ST_* values (0.39 compared to 0.14, Wilcoxon test, *P*<10^−16^), but were not any more likely to have experienced a selective sweep (*P*=0.2) than frequency-matched background SNPs (Fig. 2, Fig. SI.7-8). Absolute values of *s* were also significantly correlated with the steepness of the minor allele-environment gradients (e.g. annual precipitation [bio1] and temperature [bio12]: Spearman’s rho=0.12, *P*<10^−16^), and alleles coming from regions with low precipitation regimes tended to be positively selected in Spain under low precipitation (Fig. 2D). All in all, these observations support a polygenic model of natural selection across precipitation regimes, leading to the adaptation and specialization of local genotypes.

We finally aimed to build a quantitative environmental model that can predict *s* from past signatures of selection and local adaptation. This provided a means to predict whether alleles should increase or decrease in frequency in a certain environment, which we can use to understand evolutionary pressures in inaccessible environments or even in future hypothetical climates. We applied a regression with decision trees using Random Forests to build what we call Genome-wide Environment Selection (GWES) models (for details see Supplemental Appendix I section VII). By training models on *s* coefficients measured in Spain and Germany (10,000 random genome-wide positions), with accuracy testing of models using cross-validation and bootstrap (i.e. 10,000 other positions; 100 samples of 100), we confirmed that *s* coefficients were correctly predicted, with a high correlation accuracy (0.56 < Pearson’s r_cv_ < 0.7) and a large proportion of variance explained (R^2^_cv_= 29—52%)^24^ (Fig. 3A, <u>Table SI.7</u>). We also validated the predictability of the models using published fitness experiments with partially-overlapping sets of natural *A. thaliana* lines grown at different locations in Spain, Germany and England^25,26^ (7%<R^2^_cv_<36%, Fig. 3A, <u>Table SI.8</u>) (see further discussion in Fig. SI.11, SI.12, and Supplemental Appendix I section VIII).

**Fig. 3.**
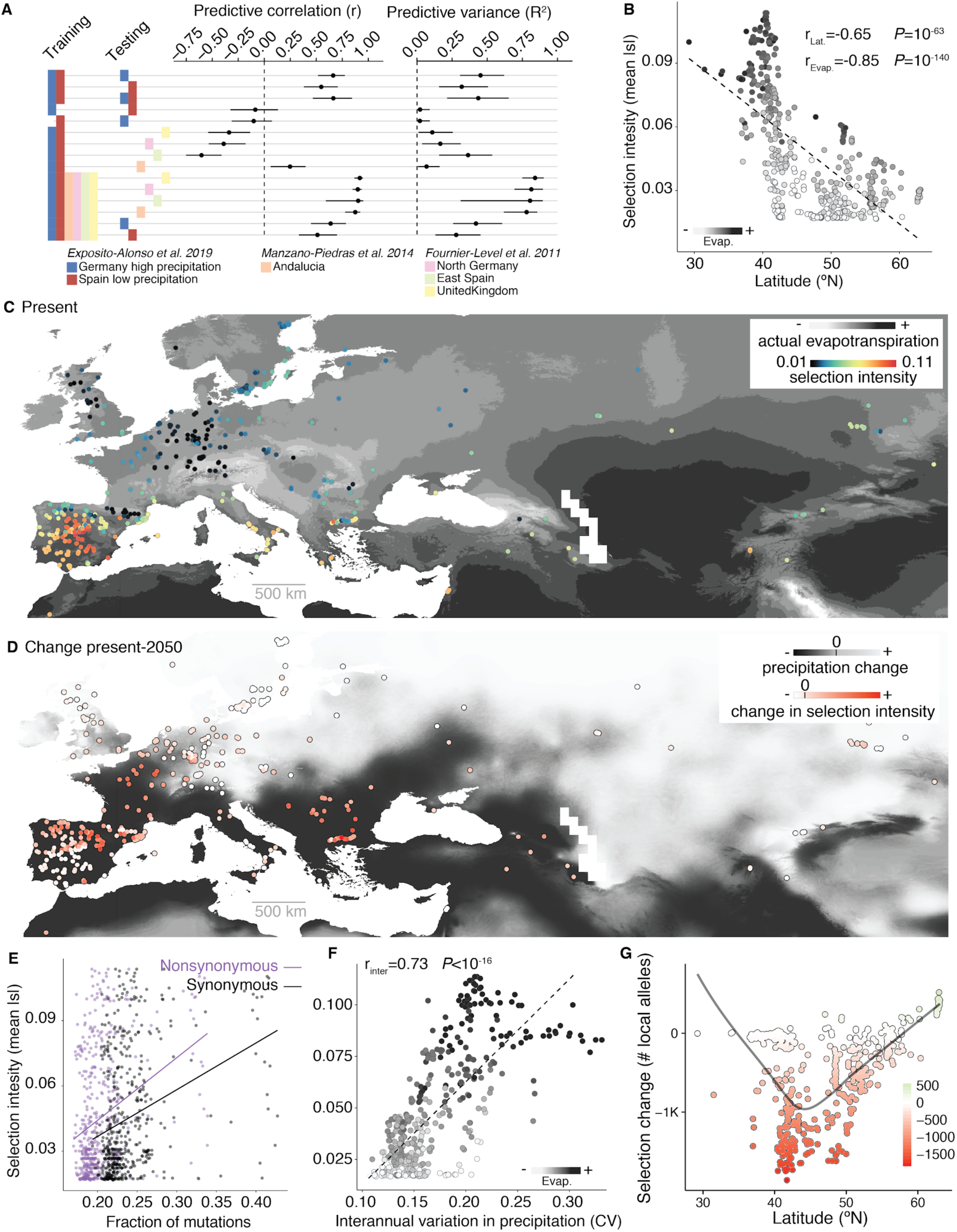
A geographic map of climate-driven selection and its predictability. (A) Genome-wide Environment Selection (GWES) models trained and tested with different combinations of our data from Germany and Spain, or from previously published field experiments (accuracy was estimated using cross-validation and the 95% confidence intervals using bootstrap, see Supplemental Appendix I section VII). (B-C) Mean GWES-predicted total selection coefficients (“selection intensity”; n= 10,752 SNPs, one random SNP per 10 kb windows) in known locations of *A. thaliana* populations in relationship to latitude and evapotranspiration in summer (ref. ^23^). (D) Predicted changes in selection intensity using climate projections to 2050 as a proxy of a sudden climate change (2050 MP rcp 8.5, ref. ^23^). (E) Relationship between selection intensity and synonymous and nonsynonymous polymorphisms present at each location. (F) Relationship between selection intensity and interannual variation in precipitation from 1958-2017 (ref. ^27^). (G) Number of local alleles (of the total 10,752 SNPs) whose selection is predicted to positively or negatively change over 5% in relative fitness by 2050 across the latitudinal range.

Using the trained and validated GWES models, we predicted genetic natural selection at hundreds of locations across the species’ native range, as if simulating field experiments in which the same set of diverse natural *A. thaliana* lines would be challenged by different local climates (Fig. 3). The intensity of selection, i.e. genome-wide average *s*, was strongest towards the environmental limits of the species, i.e. in hot (annual temperature, Spearman’s rank correlation rho=0.62, *P*<10^−16^) and dry areas (annual precipitation, rho=-0.457, *P*=10^−27^) (Fig. 3B-C, <u>Table SI.11</u>). Such environments have also a high year-to-year precipitation variability from 1958 to 2017 (rho=0.22, *P*<8×10^−7^; Fig. 3F; data from ref. ^27^). In the locations at which we predicted strong selection intensities, natural lines have a lower-than-average ratio of nonsynonymous to synonymous polymorphisms (K_n_/K_s_, Fig. 3E, rho=-0.276, 3×10^−10^, Fig. SI.14), high local genetic diversity *π*(rho=0.187, *P*=2.63×l0^−5^) and elevated Tajima’s D (rho=0.161, *P*=3×10^−4^). While several demographic scenarios could partially explain some of these patterns in isolation, they can be jointly reconciled by a scenario of strong and fluctuating climate-driven natural selection at the edge of the species range.

A sudden change in climate and increased climate variability^28^ should increase the magnitude of natural selection local population experience. Using climate projections of 2050 as a proxy for potentially abrupt changes in climate (data from ref. ^4,23^), we predict that selection intensity will likely increase in much of Southern to Central Europe, where a decrease in annual precipitation and a concomitant increase in annual temperature are expected (Fig. 3D, Fig. SI.3, SI.10).

Local populations typically consist of closely related lines that harbor only a subset of our global set of genetic variants, therefore it is important to study selection over the local variants. We therefore asked whether local alleles in each of our populations are predicted to be more or less selected in the future (change in *s* over +/- 5%) (Fig. 3G, Fig. SI.13). This revealed that many native populations in the transition zone from the Mediterranean to temperate regions will experience more negative selection, i.e. local genotypes will have a lower fitness in the future due to a diminished degree of local adaptation, putting populations at evolutionary risk.

New technological advances have enabled genome sequencing^16^ and comprehensive ecological monitoring at multiple scales^29^ for worldwide collections of plant species. Our experiments provide a proof on concept of the use of genome-wide environment selection models for evolution-aware predictions of climate change-associated risks on biodiversity. Such predictions present a first step necessary to design conservation strategies of evolutionary rescue^30^.

## ADDITIONAL INFORMATION

## Data & code availability

Phenotypic datasets are available as supplemental material with doi: [update for publication]. Genomes are available at http://1001genomes.org/data/GMI-MPI/releases/v3.1/. The seed collection can be obtained from the Arabidopsis Biological Resource Center (ABRC) under accession CS78942. The PLINK files for GWA scans of fitness and climate are deposited at aragwas.1001genomes.org with doi: https://doi.org/10.21958/study:{updatestudy_idforpublication}. Field data cleaning and processing scripts are available at github https://github.com/MoisesExpositoAlonso/dryAR with doi: 10.5281/zenodo.2583224. Plant rosette area scripts are available at http://github.com/MoisesExpositoAlonso/hippo with doi: 10.5281/zenodo.1039888 and inflorescence analysis scripts are available at http://github.com/MoisesExpositoAlonso/hitfruit with doi: 10.5281/zenodo.2583262. Simulations of total selection coefficients inference and population allele frequency changes are available at https://github.com/MoisesExpositoAlonso/selectioncorrelatedgenotypes with doi: 10.5281/zenodo.1408095.

## Author contribution

MEA, HAB and DW conceived the project outline. MEA designed, implemented and coordinated the project. MEA carried out statistical analyses with advice from RN. HAB, OB, RN, and DW supervised the project and discussed analyses interpretation. MEA prepared the first draft and the final manuscript was written by MEA, HAB, OB, RN, and DW. MEA carried out the experiment in Tübingen and in Madrid with technical support of the 500 Genomes Field Experiment Team: Moises Exposito-Alonso^1^, Rocío Gómez Rodríguez^2^, Cristina Barragán^1^, Giovanna Capovilla^1^, Eunyoung Chae^1^, Jane Devos^1^, Ezgi S. Dogan^1^, Claudia Friedemann^1^, Caspar Gross^1^, Patricia Lang^1^, Derek Lundberg^1^, Vera Middendorf^1^, Jorge Kageyama^1^, Talia Karasov^1^, Sonja Kersten^1^, Sebastian Petersen^1^, Leily Rabbani^1^, Julian Regalado^1^, Lukas Reinelt^1^, Beth Rowan^1^, Danelle K. Seymour^1^, Efthymia Symeonidi^1^, Rebecca Schwab^1^, Diep Thi Ngoc Tran^1^, Kavita Venkataramani^1^, Anna-Lena Van de Weyer^1^, François Vasseur^1^, George Wang^1^, Ronja Wedegärtner^1^, Frank Weiss^1^, Rui Wu^1^, Wanyan Xi^1^, Maricris Zaidem^1^, Wangsheng Zhu^1^, Fernando García-Arenal^2^, Hernán A. Burbano^1^, Oliver Bossdorf^3^, and Detlef Weigel^1^ (^1^Department of Molecular Biology, Max Planck Institute for Developmental Biology, Tübingen, Germany. ^2^Center for Plant Biotechnology and Genomics, Technical University of Madrid, Pozuelo de Alarcón, Spain. ^3^Institute of Ecology and Evolution, University of Tübingen, Tübingen, Germany). See author contributions to the field experiments in Supplemental Appendix II.

## Acknowledgements

We thank Patricia Lang, Angela Hancock and Talia Karasov for comments on the manuscript, and the Weigel and Burbano labs for discussions. We thank Xavi Picó for advice on experimental design, Ilja Bezrukov for advice on image processing replicability, and Belen Mendez-Vigo, Carlos Alonso-Blanco, Antolín López Quirós, Marisa López Herránz and Miguel Ángel Mora Plaza for assistance during sowing in Madrid.

## SUPPLEMENTAL APPENDIX I: Extended statistical methods for “A map of climate change-driven natural selection in *Arabidopsis thaliana*”

Moises Exposito-Alonso^1^, 500 Genomes Field Experiment Team^2^, Hernán A. Burbano^3^, Oliver Bossdorf^4^, Rasmus Nielsen^5^, Detlef Weigel^1^*

^1^Department of Molecular Biology, Max Planck Institute for Developmental Biology, 72076 Tübingen, Germany. ^2^See author contributions section. ^3^Research Group of Ancient Genomics and Evolution, Max Planck Institute for Developmental Biology, 72076 Tübingen, Germany. ^4^Institute of Evolution and Ecology, University of Tübingen, 72076 Tübingen, Germany. ^5^Departments of Integrative Biology and Statistics, University of California Berkeley, Berkeley, CA 94720, USA. Natural History Museum of Denmark, Øster Voldgade 5-7, 1350 København K, Denmark.

### I. 1001 Genomes Project data

We used VCFtools v.0.1.12b (ref. ^31^) to subset and filter the 1001 Genomes VCFv4.1 (available at: http://1001genomes.org/data/GMI-MPI/releases/v3.l/). We used vcftools with the flags: --maf 0.01 --max-alleles 2 -- min-alleles 2 --max-missing 0.95. The resulting high-quality dataset was a genome matrix of 515 individuals by 1,353,386 variants for which we did not impute the small number of missing data points.

We annotated the 1001 Genomes VCF using the package SnpEff 4.3p (ref. ^32^). We then manually curated a set of eight categories of variants: intergenic, intron, UTR3, UTR5, exon, synonymous, nonsynonymous, exon noncoding.

### II. F_st_ and selective sweep signatures from polymorphism data

We used the genetic groups previously defined for the same accessions^8^ and computed *F_ST_* using PLINK version 1.9 (ref. ^33^). We also used PLINK to calculate *π* and Tajima’s D using PLINK in windows of 100 SNPs across the genome.

We used SweepFinder2 (ref. ^34^) to scan the genome for deviations of the Site Frequency Spectrum (SFS) that might be caused by selective sweeps. We used all 11,769,920 biallelic SNPs from the 1001 Genomes Project (without the filters of 1% MAF and maximum missing data of 5%, which were applied to generate the variants used in the GWA [see section V]).

#### II.1 Geographic proxies of diversity metrics

In order to estimate *π* and a proxy of Tajima’s D at a regional scale, we used the four closest neighbouring accessions in our set (same patterns were observed with different sets of neighbours within a geographic area of 5° latitude-longitude radius), and computed the total number of polymorphisms P in the subset and the sum of all pairwise Hamming differences, H. Then we calculated *θ, π* and the proxy of D as:

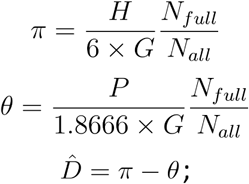

where *G* is the genome size, *N*_*full*_ are all SNPs with full information that were used to count polymorphisms and distances, and *N_all_* are all SNPs of the genome matrix. In the denominators, 6 is the number of pairwise comparisons of four genomes, and 1.8666 is the harmonic number of 4. Although D is normally divided by the standard error, we only wanted to rank our natural lines so we used the difference between *π* and *θ* as a proxy of D.

### III. Heritability of fitness

To estimate how much variance in fitness is related to the genotypes of the lines, we used generalized linear mixed models using the R package MCMCglmm with uninformative priors (ref. ^35^). We used fitness estimates per replicate and, apart from including the natural line ID, we controlled for block (growing tray) and position within the block (longitudinal, latitudinal, and the interaction). As this is a Bayesian approach, we used flat priors, we used 10,000 MCMC steps, a burn-in of 10%, and confirmed that this was sufficient for convergence of the chain. For survival proportion we used a Binomial link, for number of seeds we used a Poisson link, and for the combined lifetime relative fitness we used a Gaussian link. The mode and 95% Highest Posterior Density of the posterior distribution of each random effect were extracted (<u>Table SI.3</u>).

### IV. Interpretation of fitness GWA and consequences of natural selection for allele frequencies

#### IV.1 Total and direct selection coefficients

Genome-Wide Association (GWA) studies were first developed in the quantitative genetics field, e.g., to study common human diseases, with the main aim to determine a limited number of important loci that ultimately might have clinical or breeding utility. Because of their utilitarian motivation, these approaches focuses on specificity at the expense of sensitivity^36^. In other words, most studies focus on a few loci that can directly inform about the underlying biology.

Recently, the application of GWA approaches has been extended to other fields such as ecological and evolutionary genomics, where researchers have very different motivations, such as quantifying the total effect of natural selection at the genomic scale. For a number of organisms, it is possible to directly measure the survival of individuals or the offspring they produce. This offers the opportunity to utilize tools from quantitative genetics, where the genetic basis of a trait is interrogated, to study in addition phenomena of population genetics, where the focus is to understand population changes over time in response to selection. The latter, until now, mostly utilized indirect metrics such as allele frequency changes or low diversity islands in the genome to reconstruct the fitness consequences of mutations or their proximity to fitness-related variants.

The potential of utilizing GWA with fitness traits to study population genetic phenomena was best described in a recent paper from Gompert and colleagues^17^. They used statistical software developed for phenotypic GWA and thoroughly discussed the differences in interpretation of multiple SNP-fitness association estimates in this context, contrasting them with phenotypic GWA effects. Specifically, GWA techniques allow for calculating <u>total</u> and <u>direct</u> effects of natural selection over SNPs. To illustrate the point here, we consider the most simple case of two biallelic SNPs, *x*_1_ and *x*_2_, with three possible genotypes in a diploid individual *x_i_* ∈ {0,1,2}. This can later be extended to large numbers of SNPs, something that has been widely addressed by GWA methods^18^. For mathematical convenience we assume that the predictors (*x*_1_ and *x*_2_) are mean centered and variance scaled, while the response variable fitness, *y* is relative fitness. From the univariate approach, where the effect of a SNP *x*_1_ is estimated <u>marginally or independently</u> from *x*_2_, the <u>total effect in selection</u> would be:

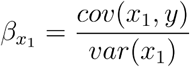

The same calculation would be repeated for SNP *x*_2_. In a multivariate regression framework, the regression coefficient, called <u>conditional or partial</u> coefficient, *β*^*^, is corrected by the correlative indirect effect of the other predictor, *r*_*x*_1_*x*_2__. In this way, effects driven by linkage disequilibrium to other loci are removed from *β*\*, so that only <u>direct effects</u> are measured. The formula would be as:

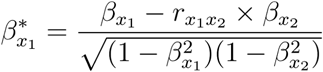

*β* would capture the total selection and thus can be called a *“total selection coefficient”*, as *β_x1_* ≈*s* = *w*_*x*1_=22 − *w*_*x*1_=00, where *w* represents the relative fitness of a plant genotype. On the other hand, *β\** cancels out indirect (or linked) effects, and thus can be called *“direct selection coefficient”*.

Gompert and colleagues^17^ use the statistical GWA package GEMMA (ref. ^18^) to infer the above coefficients for thousands to millions of SNPs. GEMMA implements a single-marker <u>marginal linear models GWA</u> (LM) of the form: y = *μ* + *β_i_x_i_* +*ϵ*; which provided us with each allele’s marginal effect *β* on relative fitness per SNP, or total selection coefficient. GEMMA implements a Bayesian Sparse Linear Mixed model GWA (BSLMM), which allows the genome-wide calculation of direct (*β^*^*)selection coefficients (e.g. n=1,353,386). GWA models have been specifically developed to solve the problem of the number of predictors, in this case SNPs, vastly exceeding the number of observations, such as trait or fitness values. These models come in two related flavours: (1) the random-genotype-effect-with-known-relatedness matrix (also called kinship or GBLUP GWA). which captures the joint correlation of many SNPs across genotypes. (2) the joint fitting of all SNPs, as in our two-SNP example, but shrinking the effects (e.g. using ridge, lasso, or shrinking priors approaches). BSLMM does both at the same time. It models two effect hyperparameters per SNP, a basal effect *α* that captures the fact that many SNPs contribute to the phenotype, and an extra effect *β* that captures the stronger effect of only a subset of SNPs. An internal parameter measuring the probability of having an extra effect *γ* can be used to prioritize SNPs. In BSLMM the overall effect of an allele is = *α* +*γβ*, which in the simple example of two SNPs above would correspond to *β*.* The full model specification is:

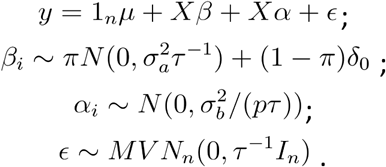

Because this model corrects for correlated effects among SNPs, it effectively corrects indirect linked (LD) effects arising from population structure and/or low recombination. Both BSLMM and GBLUP GWA approaches enable one to calculate the total proportion of variance explained by genetics (PVE or “chip heritability”). In our analyses, we used the last 1,000 samples of the MCMC chain in BSLMM to calculate the median and 95% Highest Posterior Density Interval (95% HPD). As mentioned above, the BSLMM model is related to the classic GBLUP or kinship-based GWA, a form of linear mixed model where one corrects-out population structure or general relatedness between individuals. For completeness, this is done by having the *u* random effect term with a given covariance structure, that is, the kinship matrix *K*, and the calculated *β^*^* is conditioned on genomic background effects:

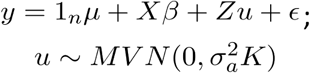

The two estimates above, the direct (conditional *β\**) effect calculated from BSLMM or GBLUP GWA and the total (*β* or *s*) effect calculated from marginal GWA, provide thus different insights on the nature of selection, which are both useful in their own right. As already argued by Gompert and colleagues^17^ as well as others, it is the total selection coefficient *s* that best predicts the change in population allele frequency in one generation as a response to selection. We show this with simulations in the next section.

#### IV.2 Proof-of-concept simulations on the importance of total selection coefficients

To illustrate the prediction accuracy of frequency changes in a population from total selection coefficients ***s*** or direct selection coefficients, we carried out simulations (Fig. SI.15) (see code: https://github.com/MoisesExpositoAlonso/selectioncorrelatedgenotvpes)•

We began by subsetting our dataset of 515 genomes of *A. thaliana* to 1,000 consecutive SNPs from chromosome 1 (to reduce the complexity of calculations while keeping the linkage structure intact. Note: Our results also hold when simulating a genome matrix with random linkage disequilibrium). We then assigned 1,000 selection coefficients (i.e. true direct selection coefficients) to the 1,000 SNPs following a normal distribution with mean zero and standard deviation 0.1. To obtain the fitness of a plant of genotype *j*, we sum selection coefficients along the genome as: 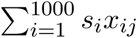 where *x*_*ij*_ indicates whether the haplotype has the reference (00) or alternative allele (11) in the given *i*th SNP (note in this *A. thaliana* dataset there are no heterozygotes, as these are naturally inbred lines that have been further selfed in the laboratory). For realism, we added some artificial noise to the plant fitness (heritability=0.9, although conclusions hold with other intermediate heritabilities). We then inferred <u>total selection coefficients</u> using marginal GWA and <u>direct selection coefficients</u> using GBLUP GWAs. Our results comparing true and estimated effects show how we largely fail even when we attempt to estimate <u>true selection coefficients</u> with the GBLUP method, which tries to correct by background effects (Fig. SI.15A). It is also important to notice that the estimates from GBLUP GWA are one order of magnitude smaller than the true values. Because the architecture of fitness here is rather polygenic (as in our experiments) and SNPs are in linkage, most of the fitness variation is assigned to the kinship term in the GBLUP GWA rather than to specific SNPs. In agreement, the kinship-based random effect accumulates 99% of the true heritability (*Vg*/*Vg* + *Ve* = 0.89). In our field experiment, we also saw high values of kinship-based heritability (our median h^2^ was 0.7; <u>Table SI.2</u>).

As discussed earlier, if the interest is on the <u>consequences of natural selection in genome-wide on allele frequencies</u> rather than identifying only the alleles under direct selection, estimating the <u>total selection coefficient</u> is the most adequate approach. To show this, we ran *in silico* an individual-based simulation, reproducing genotypes proportionally to their relative fitness to generate the population of offspring one generation after selection (with constant population size). We then compared the change in frequency in the simulated population with the marginal GWA and GBLUP GWA estimates. Because <u>frequency changes at a given allele are driven both by direct and indirect selection</u> pressures, the marginal GWA estimates correlate best with the changes of frequency in one generation (R^2^=0.39 Fig. SI.15B, compared to GBLUP-based R^2^=0.02). In fact, we can try to predict directly the change in frequency in one generation (Δ*q*) if we not only use inferred selection coefficients but also the original starting frequency, as the effect of selection is also proportional to the starting frequency of an alleles: Δ*q* = *S*(1 − *p*)*p.* plugging in the marginal GWA and GBLUP GWA estimates into the equation, we show that marginal GWA estimates allow prediction of allele frequency change with accuracy of R^2^=0.97, while GBLUP GWA estimates are extremely poor, R^2^=0.08 (Fig. SI.15C). Of course, this is again because of the influence of linked effects of selection.

A potential concern of studying total selection coefficients rather than direct selection coefficients is that the first are thought to be completely contingent on the specific allele frequency and linkage structure of the population analyzed. In practice, in many cases allele frequencies and linkage structures will not vary sufficiently to invalidate any extrapolation. Our experimental population of 515 accessions (subset of the 1001 Genomes Project) aimed to maximize geographic as well as genetic coverage of the species, which in turn enables interpolations to smaller, less diverse regional subpopulations.

We then studied to what degree measurements of natural selection hold across populations. Out of the global set of 515 *A. thaliana* lines, we selected 50 Spanish genotypes, which are known to belong to very distinct lineages. We run individual-based simulations with the 50 Spanish genotypes to compute allele frequency changes. We used the marginal and GBLUP GWA estimates calculated with the 515 genomes to predict allele frequency changes and correlated the results with the simulations (Fig. SI.15D). With this we could show that marginal GWA effects calculated in the global panel of 515 lines also predict well the frequency changes after selection in the subset of 50 Spanish lines (R^2^=0.85), and much better than the GBLUP GWA (R^2^=0.04). (NB1: we validated these conclusion with other subset populations, such as 10 Spanish accessions or 10 US accessions with low genetic diversity. NB2: The converse was also true, namely that marginal GWA effects calculated in the 50 Spanish lines predicted well allele frequency changes in the global panel of 515 lines, R^2^=0.76).

#### IV.3 Further notes on the interpretation of population structure in wild species

Much of the discussion in the previous section was about the special interpretation of marginal GWA estimates on relative fitness as total selection coefficients, which have the property of predicting allele frequency changes. We stress that relative fitness is a very special trait. It is exceptional in terms of the extra insight that can be gained from a marginal GWA, which is very different from other traits, where the estimates do not have any association with total selection coefficients nor allele frequency changes. For GWAs of morphological or physiological traits, there is no dispute that population structure-or linkage disequilibrium-aware GWA approaches such as GBLUP or BSLMM are the best choice (e.g. refs. ^8,37^).

In this context, it is worth recalling why population structure correction in human GWA studies addresses issues that are not necessarily relevant for wild species that can be grown in common gardens: Human GWA studies are typically based on data collected in different countries or localities, where individuals were born and raised in different, often country-specific environments. Because such factors are correlated with spatial location and genetic ancestry of human populations, the population structure correction also corrects for such structured environmental noises. In experimentally-tractable wild species, such confounders are easily avoided by controlled experiments and replication. In cases where phenotypic data of wild species come directly from field observations, as in humans, population structure correction remains paramount.

As a final note: in wild species, there might be cases when population history and differentiation coincide with historical events of adaptation. Here, population structure correction can make it challenging to correctly identify valuable SNPs, as most of the true positive SNPs will be tightly correlated with historic population lineages and thus much (if not most) of the important variation will be assigned to the kinship factor, relegating these valuable SNPs to no-effect SNPs (false negatives). In such cases, it is up to the researcher to decide what is the best method in a case-specific manner. Some approaches such as the BSLMM approach try to solve background effect confounders more elegantly than the kinship approaches, but they have other limitations^17^. Another potentially useful approach is admixture mapping^8,38^.

This section does not aim to de-legitimize population correction approaches for wild species nor can it serve as a comprehensive discussion on the topic. Instead, it hopes to open up a discussion of potential new uses of GWAs and be thought-provoking to ensure researchers carefully design and interpret their ecological genomic analyses.

#### IV.4 Trade-offs of selection

##### IV.1.1 Across field experiments

In order to test the two most prevalent hypothesis of local adaptation driven by selection trade offs, conditional neutrality vs antagonistic pleiotropy^22,25,39^, we do pairwise comparisons of total selection coefficients in two environments. We devised two tests: The first test discriminates between pleiotropy (selected either in the same direction or in different directions) and conditional neutrality (only selected in one environment). We use the extreme 5% selection coefficients at each tail, similar following Anderson et al. ^39^ to generate the contingency table:

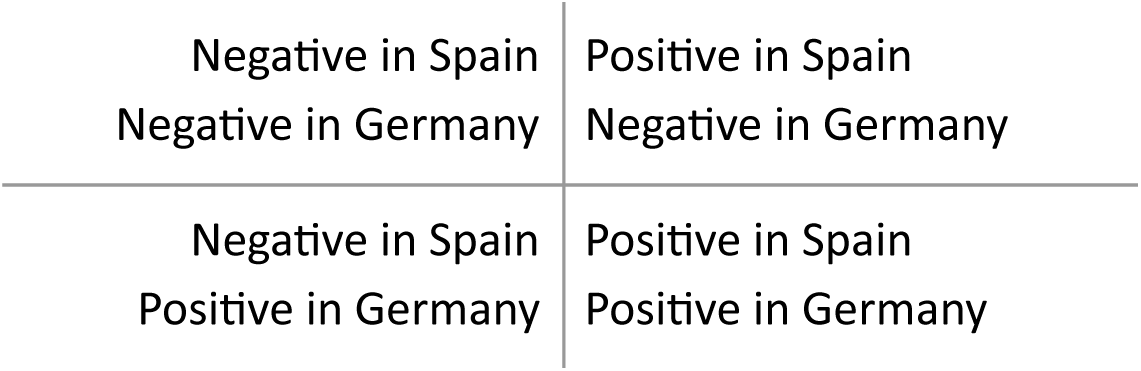

Because this test does not distinguish between those pleiotropic variants that are selected in opposite directions (antagonistic pleiotropy) or in the same direction (non-antagonistic or synergistic pleiotropy), we do another test only for the direction of those variants selected in both environments:

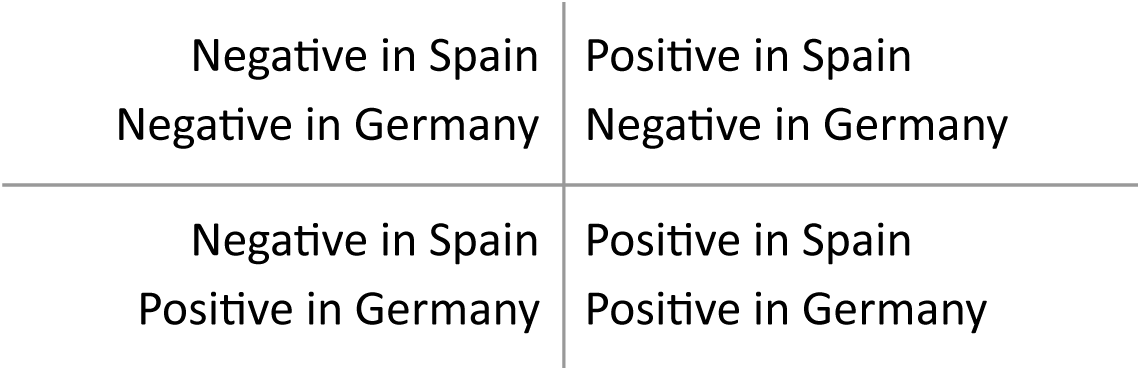

We report the Odds Ratio for both tests (<u>Table SI.5</u>) as well as the Spearman’s rho correlation between each pair of environments (<u>Table SI.6</u>). The tests in (<u>Table SI.5</u>) were conducted with all SNPs (N=1,353,386), but in order to check that the lack of independence did not affect the Odds Ratio, we re-calculated our analyses using one random SNP per 100Kb window, with similar result (e.g. comparison Spain low precipitation high density vs Germany high precipitation low density, Odds pleiotropy vs conditional neutrality = 3.0419, *P*=2.627×10-5; Odds antagonistic vs synergistic pleiotropy=15.979, P=0.0093).

##### IV.1.1 Across life history stages

Calculating total selection coefficients for survival and fecundity separately, we found no correlation between survival-only and fecundity-only estimates (r<0.07, Fig. SI.4-5), consistent with different stages of a plant being differentially affected by environmentally imposed selection^40,41^.

#### IV.5 Intensity of selection

The distribution of absolute total selection coefficients, |*S*|, has a shape resembling that of an exponential function. We calculated the expected rate using Maximum Likelihood optimization in R, which can also can be approximated as the inverse of the mean:

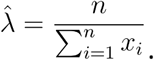

For this, we use 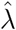 or the mean of |*S*| as a metric of the overall intensity of selection (Fig. 1B, Fig. 3D).

### V. Climate Genome-Wide Association

Similarly to our GWA with relative fitness, we run a GWA with each climate variable *m* (see Section VII.1) as response variable *y_m_* in a LM model using GEMMA (ref.^18^, see Section V): *y*_*m*_ = *μ* + *β*_*i*_*x*_*i*_ + *ϵ*;

This *β* coefficient for SNP ***i***, which reflects the correlation of the alternative allele’s presence and a climate variable, was used later in our predictive models (<u>Section VIII).</u> As this is a raw correlation between allele presence and climate variables, it will capture both past signatures of climate adaptation and historic population migration and differentiation, and is only used to capture how environmentally separated are typically found the two alleles of a SNP. Because the purpose was to capture climate associations that reflect past adaptation, we only used natural lines from within the native range, within the 25 to 65° Latitude and −15 to 100° Longitude (502 native lines of the total 515).

The top 1% hits for climate associations of multiple variables had higher *F_ST_* values than frequency-matched background SNPs (e.g. bio1 and bio12: *P*<10^−5^), but no differences in sweep likelihood (*P*=0.9).

### VI. Climate and modeling

#### VI.1. Climate layers

We used the classic bioclim variables (n=19), plus monthly data of minimum and maximum temperature, and precipitation (n=12 × 3) (worldclim.org). From these we estimated monthly evapotranspiration rates using the R package EcoHydRology v. 0.4.12 (ref. ^42^) and actual monthly evapotranspiration using a bucket model ^43^ (n=12 × 2). Based on ref. ^44^ we calculated whether *A. thaliana* can grow in a given month based on temperature and precipitation (n=12), and derived from this the length of the potential growing season (n=1). Over the potential growing season, we calculated minimum and maximum temperature, and total precipitation (n=3). Finally, using the mean and variance flowering time (=lifespan) across all our field experiments per accession, and based on their climate of origin using the above variables, we used an environmental niche model to generate a map surface of the most likely plant lifespan (n=2). This provides an estimate of the actual growing season, which we subtracted from the potential growing season to generate one more composite variable (n=1). Each variable is further described in <u>Table SI.9.</u> A total of 98 raster layers are available as .gri/.grd files (native R format) from: github.com/MoisesExpositoAlonso/araenv.

#### VI.2. Environmental Niche Models

Genome-wide Environmental Niche Models (GEMs) were fit using decision trees with presence/absence of SNPs as response variable and the climate variables described in the previous section and latitude and longitude as predictors as described previously^8^. To fit the models, we used a Stochastic Gradient Boosting approach with the R package caret (ref. ^45^). The parameters used to fit the model were: 50 decision trees, an interaction depth of 2, a shrinkage of 0.1, and a minimum of observations at end nodes of 10. This set of parameters was determined after running our GEMs for some exemplary SNPs and confirming that this set was typically optimal for reducing residual-mean squared error in a Repeated Cross-Validation approach.

We used these models to predict from raster maps of the climate layers a probability between 0 and 1 that the alternative allele was in a map cell. We judge this as a more appropriate output than a discrete 0/1 outcomes, as sometimes alleles were widespread or at intermediate frequencies in many regions and thus their environment niche was not strictly defined.

Areas outside the high-density areas of *A*. *thaliana* (Fig SI.1) were excluded from the GEM training and projections, as our information of populations, for instance, from Siberia is limited. Nevertheless, the few samples there had a relatively high fitness in Spain and low precipitation (<u>Dataset 3).</u>

#### VI.3. Climate variability

To study spatial climate variability, for each *A. thaliana* natural line, we extracted the 19 bioclim variables (<u>Table SI.9</u>) in a 50 Km buffer where they were originally collected from and calculated the coefficient of variation (CV) across grid cells.

To study temporal variability, we used climate data^27^ from 1958-2017 to calculate annual precipitation values for each population, from which we in turn derived the inter-annual CV.

### VII. Predictions of total selection coefficients from sequence and climate features

#### VII.1 The model

We used a decision tree approach with Random Forest using the R package randomForest (ref. ^46,47^) to predict the vector (n=1,353,386) of GWA results with relative fitness in one environment, which we call total selection coefficients *s*, from a 1,353,386 × 98 matrix of GWA associations with climate variables, *β_dim_*(<u>Table SI.9</u>, section V). We also included as predictors a 1,353,386 × 5 matrix, *μ*, of genetic diversity and frequency metrics: minimum allele frequency, π diversity, Tajima’s D, selective sweep likelihood ratio, and selective sweep alpha value (section II). In addition, we included as predictors a 1,353,386 × 8 matrix *θ* of non-mutually exclusive variables taking values of 0 or 1 indicating genomic annotations: intergenic, intron, UTR3, UTR5, exon, synonymous, nonsynonymous, exon noncoding (section I). A total of 112 variables were thus used as predictors:*s* =*f*(*β_dim_,μ,θ*). In the cases where we trained models with two environments, we also included the 2 × 98 *x_dim_* climate variables at our field stations: *s* =*f*(*x_dim_*,*β_dim_,μ,θ*).

Conceptually, GWES models are similar to Environmental Niche Models (ENMs), but instead of training them with presence/absence data of a genetic variant^8^, we trained them with our measured total selection coefficients.

#### VII.2 Genome-wide cross-validation

Because training a Random Forest with the full dataset would be computationally expensive, we only trained with 10,000 observations (with smaller and larger SNP sets, we had determined that training with more than 10,000 observations did not improve predictions). To test accuracy and bias we used a different set of 10,000 SNPs, divided into 100 bootstrap samples, and we report the intervals of the 95% bootstrap distribution. The results presented in Fig. 3 were produced with 10,000 randomly drawn SNPs across the genome. To confirm that there was no confounding from non-independent samples in the training and testing SNPs, we repeated all analyses, training with 10,000 random SNPs from chromosome 1 and testing with 10,000 random SNPs from the four other chromosomes. There were no substantial changes in predictability.

Several combinations of training and testing were performed to validate the predictions of “unobserved” environments (<u>Table SI.8).</u>

#### VII.3 Note on interpretations and limitations of GWES

> *“It’s Difficult to Make Predictions, Especially About the Future.”*
>
> *(unknown, but often attributed to Yogi Berra)*

As in any predictive exercise, our geographic projections of intensity of selection have limitations. We believe they are nevertheless indispensable to move forward in the field of forecasting climate impacts. Models such as ours are tremendously useful for subsequent experimental validation (as we are currently doing through an experimental evolution network: GrENE-net.org) or with *in situ* observations collected as we move into the future (e.g. iNaturalist.org, iSpot.org). This iterative prediction ↔ validation process will be key to advancing the complex field of predicting the effects of climate change on biodiversity.

To make detailed predictions for specific local populations in 2050 will require knowledge about local genetic diversity, population sizes, migration in meta-populations, outcrossing rates, reliable year-to-year climate stochasticity models, and any other potential demographic and environmental factor that affects evolvability. Our measured selection coefficients will help to build such models. In the meantime, our predictions serve as important indicators of relative evolutionary risk that local *A. thaliana* populations are likely to experience in the near future.

Below we discuss potential pitfalls of GWES, and the Dos and Don’ts of GWES interpretation.

A. Selection is a “relative force”. The selection of a specific allele depends on the alternative allele, and at what frequency both are found. Thus, the exact value of total selection coefficients might vary depending on the GWA panel. A *reductio ad absurdum* case would be that total selection coefficients are estimated in a diverse GWA panel, but they are invariant in a specific population. In such a case, one could not calculate total selection coefficients for invariant sites in this population, although that does not mean this population is not under natural selection, which could lead to its extinction if it is fixed for the disadvantageous alleles. As we discuss in Section IV.2, and show with simulations in Fig. SI.15D, by using a diverse reference GWA panel to calculate total selection coefficients, we can interpolate to subset populations. Therefore the GWES projections are useful for relative trends of selection across the species’ geographic range.
B. Our GWES projections are not long-term population projections, which need to take into account many other factors:

a. Short-term total selection coefficients (over one generation, ecological times) do not necessarily reflect long-term selection coefficients (i.e. over evolutionary times), which are an integration of selection events over time.
b. Over longer timescales, immigration of genotypes, admixture, and recombination, can alter the efficiency of selection.
c. Demographic dynamics are ultimately determined both by natural selection and stochastic demographic forces (drift). Therefore, the knowledge of total selection coefficients in one generation is necessary, but not sufficient to determine the fate of a population over multiple generations. To do so, explicit demographic models are needed, models that’ also take into account nuances such as bet-hedging strategies with seedbanks, and overlapping generations.
d. We fed climate projections of 2050 (of different CO_2_ scenarios) into the GWES models as proxies of plausible magnitudes of short-term climate changes. Year-to-year demographic processes will interact with the local gradual or stochastic changes in climate and ultimately determine the extinction or persistence of populations. A useful way to think of our climate projections from models trained at the warm edge (Spain) and in the distribution center (Germany) is to think how climate change might put German populations under selection pressures similar to those prevailing in Spain.

### VIII. Re-analysis of published data from common garden experiments

#### VIII.1 Environment cross-validation

In order to cross-validate our model on independent environments, we re-analyzed published data. This approach is an environmental cross-validation on top of cross-validation of SNPs. That is, we train in a subset of 10,000 SNPs in Spain and Germany, and test our model in another subset of 10,000 SNPs using the previously-published experiments of Spain, Germany and England^25,26^. For a conceptual diagram of predictability (and extrapolability) validation with external common garden experimental datasets, see Fig. SI.11. Note that a partial overlap of natural lines and genomic data is required for the following re-analysis (predictions on common gardens with recombinant inbred lines or non-overlapping natural lines would require further adjustments in our approach).

#### VIII.2 Manzano-Piedras et al. 2014

Manzano-Piedras and colleagues^26^ planted exactly 60 seeds per line in pots. They monitored how many plants established at the rosette stage and later on became reproductive adults (survival proportion). From these, they counted the number of fruits per pot and divided them by the number of reproductive adults (reproduction, seed set). We computed lifetime fitness as the product of survival and reproduction.

#### VIII.3 Fournier-Level et al. 2011

Fournier-Level and colleagues^25^ germinated seeds in greenhouses, and two weeks after germination (established seedling stage), they transplanted seedlings to outdoor field stations where one plant was transplanted in one pot. They counted how many transplanted seedlings survived to reproduction (partial survival proportion), and the number of fruits per plant (reproduction, seed set). We again computed lifetime fitness as the product of partial survival and reproduction.

We excluded the experiment in Finland in downstream analyses because only 58 natural lines were planted there in the original publication^25^ and because we found the imputation accuracy to be very low (Pearson’s r<0.008).

#### VIII.4 1001 Genomes x RegMap panel phenotype imputation

The 1001 Genomes panel (http://1001genomes.org/, ref. ^16^) includes 1,135 natural lines with 11,769,222 biallelic SNPs from Illumina sequencing. The RegMap panel (http://arabidopsis.gmi.oeaw.ac.at:5000/DisplavResults, ref. ^48^) includes 1,307 natural lines with 214,051 biallelic SNPs from array hybridization. The two populations shared 413 lines. Of these, 185 were shared with the 515 lines used in the field experiments.

Of the 157 accessions of Fournier-Level *et al.*, all were part of the RegMap panel, 89 were part of the 1001 Genomes, and 50 overlapped with our lines. Of the 279 accessions of Manzano-Piedras *et al.*, 150 were part of the 1001 Genomes, and 131 overlapped with our field lines.

Because fitness is heritable, we tried to impute missing data based on the overall genomic relationships among all of the 2,029 natural lines belonging to 1001 Genomes and RegMap panels. After downloading and transforming the RegMap dataset to PLINK format, we overlapped genome-wide SNPs and filtered them for a genotyping rate of 95%, which yielded 154,090 biallelic SNPs. Given the linkage disequilibrium and genome size of *A. thaliana*, this easily suffices for generating a relationship matrix *A* (related to a kinship matrix), which we computed using the R package rrBLUP (ref. ^49^). The data of survival, reproduction, and lifetime fitness was an average per genotype, so we fit a classic GBLUP: *y = Zg + ϵ* where *y* is the fitness trait of interest, is a design matrix of genotypes and *g* is a random effect factor with covariance matrix equal to the relationship matrix 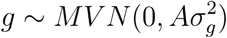. Heritability of traits and imputation accuracy from the Manzano *et al.* and Fournier-Level *et al.* experiments is given in <u>Table SI.10.</u>

#### VIII.5 Sanity checks for imputation and geographic predictions

We carried out sanity checks to ensure that the imputed fitness from other experiments was not just an artifactual phenotype with the same structure as the relationship matrix. This would mislead us to find predictability even without causality, as we would expect that total selection coefficient calculated in such artifactual phenotype would depend on population structure and thus would likely be predictable from climate structure alone.

We shuffled the genotype identities from Fournier-Level *et al.* and Manzano-Piedras *et al.* with their fitness values. Then we repeated the GBLUP analysis with 50 rounds of shuffling and computed heritabilities and prediction accuracies. We confirmed that heritability with shuffled data was negligible (1xl0^−9^<h^2^<1.6^−3^) and so was the accuracy of imputation (−0.047< r <0.070). This indicated that in the absence of true heritable variation, imputation of fitness would be random and not an artifact of the relationship matrix.

We also were concerned that geographic predictions could be driven by some underlying bias in our analyses, i.e. bias inherent to geographic sampling, population history of genotypes chosen, etc. In other words, we were concerned that the null expectation of predictability would be non-zero. As before, we randomized fitness values with genotypes for all six environments (Fournier-Level *et al.*, Manzano-Piedras *et al.*, and ours). Then, we repeated the GWA to estimate total selection coefficients (as Fig. 1), and trained different combinations of GWES models to re-predict total selection coefficients at each location based on climate (as Fig. 3). We confirmed that, differently from the analyses of real data presented in Fig. 3, there was no significant predictability (Fig SI.12).

#### VIII.6 An explanation for “inverse predictability”

We noticed that using only our two experiments for model training, there was “inverse predictability” for the three experiments from ref. ^25^. While the sign of inferred total selection coefficients was the opposite of the observed values (−0.33<r_cv_<-0.51, *P*<0.001), the magnitude of selection was correctly inferred (15%<R^2^_CV_<25%, Fig. 3A). Such a phenomenon could arise for several reasons and has already been observed in studies with evolving *Drosophila* populations where seasonal environments vary from year to year^50^, as well as in timema insects^24^. In our case, the worldclim.org climate averages (1960-1990) at 2.5 arc-minutes resolution might strongly deviate from the truly experienced environmental conditions in the years the experiments were conducted. Such climate variability can exert opposite selection in different years^51^. Second, differences in experimental design could lead to different lifetime fitness estimates. In the Fournier-level *et al.* experiments, early survival of seedlings was not measured at all, as only seedlings that had survived for two weeks in the greenhouse were transplanted into the field. In the Southern Spain experiment from Manzano-Piedras *et al.* ^26^, seeds were sown directly in the field, as in our own experiments, and accordingly, we had “positive predictability” (r=0.24, Bootstrap Cl=0.09—0.41). In further support of this experimental design confounder, when we trained GWES models with only reproduction-based total selection coefficients in our experiment of high precipitation in Southern Germany, i.e., excluding early survival from lifetime fitness, we correctly predicted the sign of total selection coefficients in Fournier’s Northern Germany experiment (r=0.392, Bootstrap CI=0.20—0.57) (for null expectations see <u>Supplemental Methods IX.4</u>).

The differences in predictions between two-and six-environment-trained models did not yield differences in downstream conclusions from Fig. 3 (correlation between predictions, r=0.56, *P*<10^−16^), but predictability increased with the number of experiments included in the training set (6 environments, r= 0.746 [Bootstrap CI= 0.667— 0.800], R^2^= 0.517 [0.445–0.640]).

We preferred to show geographic predictions (Fig. 3) with GWES trained with our two environments so we only rely on highly-replicated fitness estimates from over 500 accessions that were grown in carefully controlled precipitation and temperature environments.

---

**Figure SI.1.**
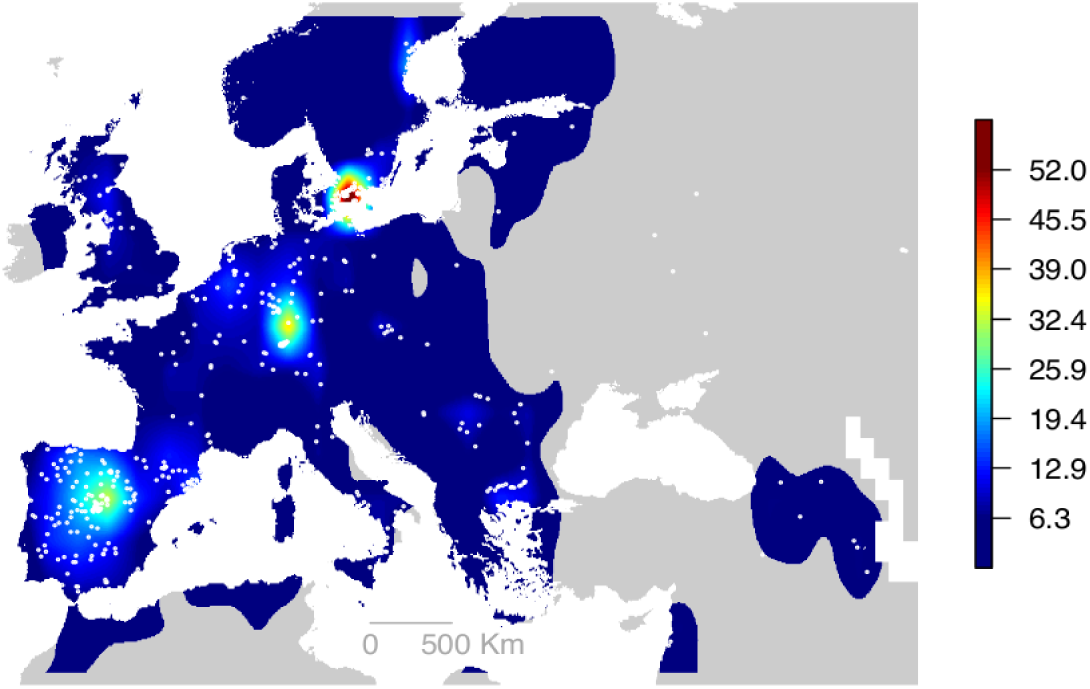
Map of abundance of *Arabidopsis*. samples Points indicate the locations where the 517 *A. thaliana* accessions were collected. The color gradient is the density of samples from our study in squares of approximately 200 km × 200 km. The limits of the colored area were determined using a combined density grid from gbif.org and 1001 Genomes records. The density was generated in a grid of 125 min resolution and by applying a bilinear and then Gaussian smoothing. The threshold was chosen to be the 50% of the upper distribution, which roughly corresponds to 10 records per 200 km × 200 km square. Regions outside the colored areas were excluded in allele’s environmental niche models (Fig. 1), as we prefer to make predictions only in regions where the presence of *A. thaliana* is rather likely and continuous. Predictions of future climate change-driven natural selection (Fig. 3) were taken here with caution for the same reason.

**Figure SI.2.**
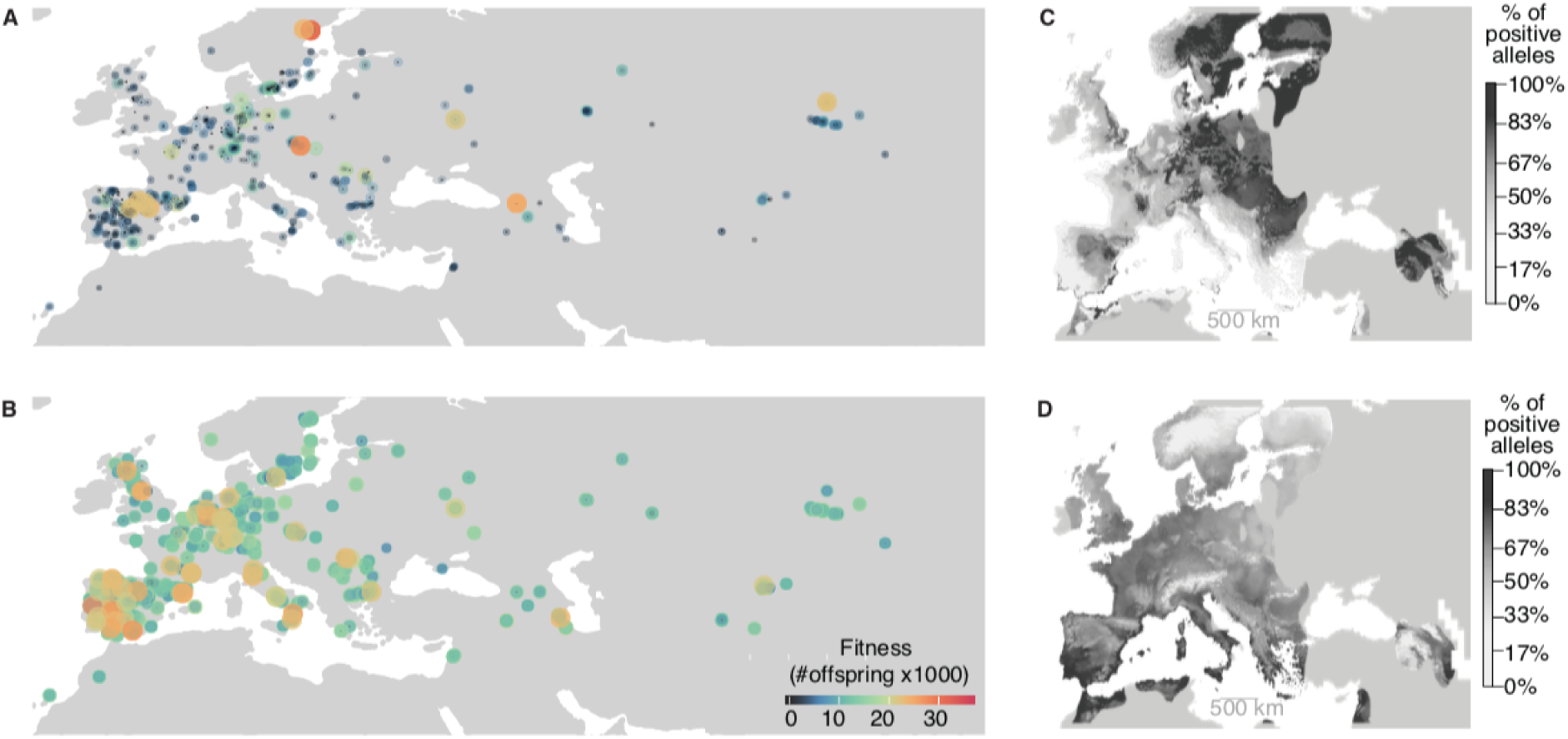
Environment ranges. (A, B) Raw fitness data (# offspring produced by each pot), in Madrid low precipitation (A) and Germany high precipitation (B). In agreement with previous observations of survival of terminal drought treatments in the greenhouse^8^, the most successful genotypes in Madrid and low precipitation come from Spain, but also other areas of the distribution with extreme climate, including North Sweden, Eastern Europe, the Caucasus, and Siberia. While in Germany and high precipitation there seems to be a trend that lower latitude genotypes, this could also be just a reflection that in the Mediterranean there are more diverse genotypes (some with higher and some with lower fitness than average). All in all, variation in fitness in Germany is very low and less genetically heritable (<u>Table SI.3</u>). (C, D) Idealized representation of alleles distributions identified in Spain and Germany as inferred from Genome-wide Environmental Niche Models. The most significant SNPs in each 0.5 Mb window of the genome was modeled. Color scale indicates the % of locally present alleles with respect to the maximum number of positive alleles. (C) 424 genomic windows had significant SNPs in high-precipitation experiments. (D) 279 windows had significant SNPs in low-precipitation experiments. The maps of alleles generally agree with the trends observed in (A,B).

**Figure SI.3.**
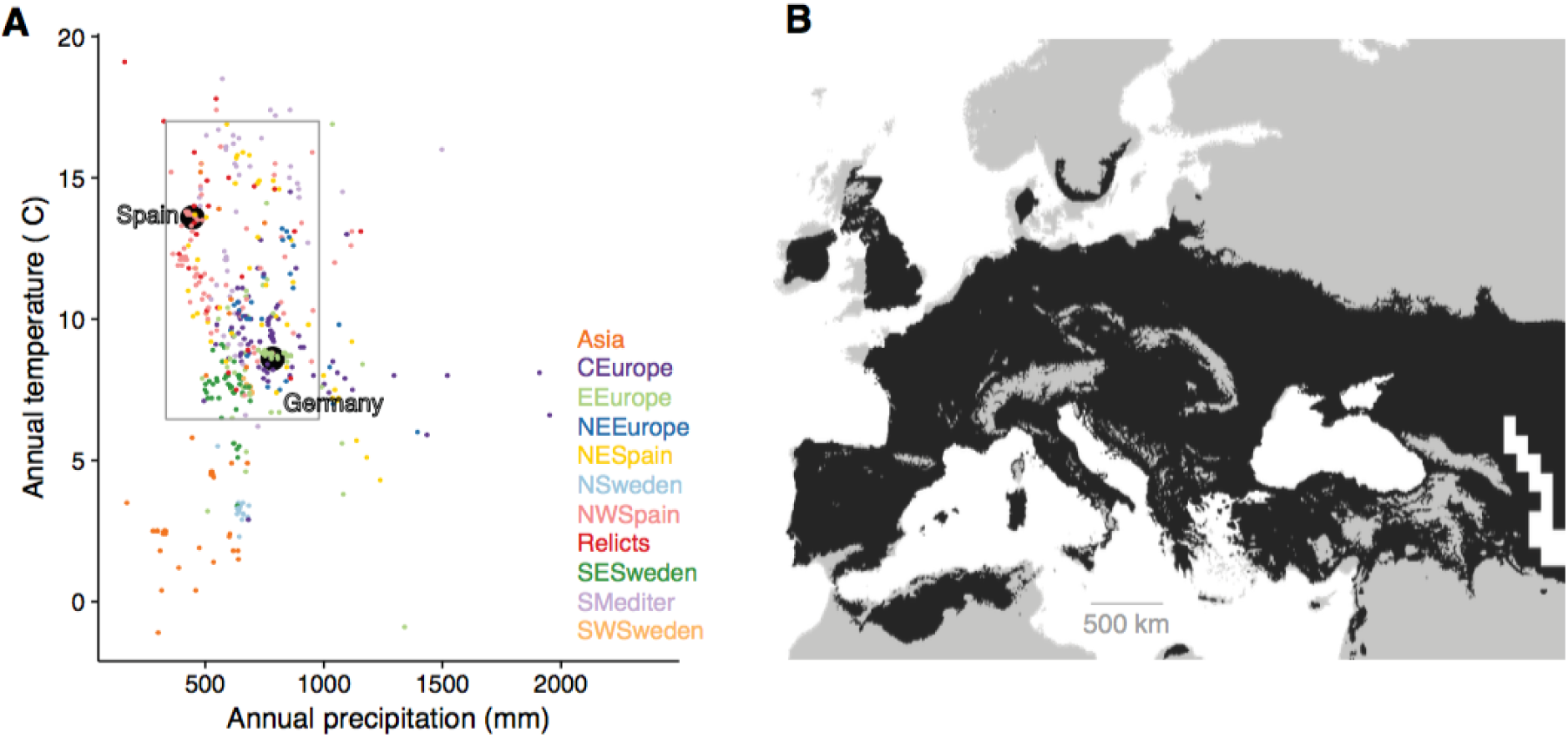
Map of predicted precipitation change. (A) Classic biplot of precipitation *vs.* temperature of origin of accessions (black dots) and field experiment of Spain (sepia) and Germany (green). Grey box indicates locations where precipitation was at least 70% of Spain and no more than 130% of Germany, and where temperature was no less than 70% of Germany and no more than 130% of Spain. (B) Areas that would be within the grey box in (A). Colored population groups based on previously calculated genetic clusters^8^.

**Figure SI.4.**
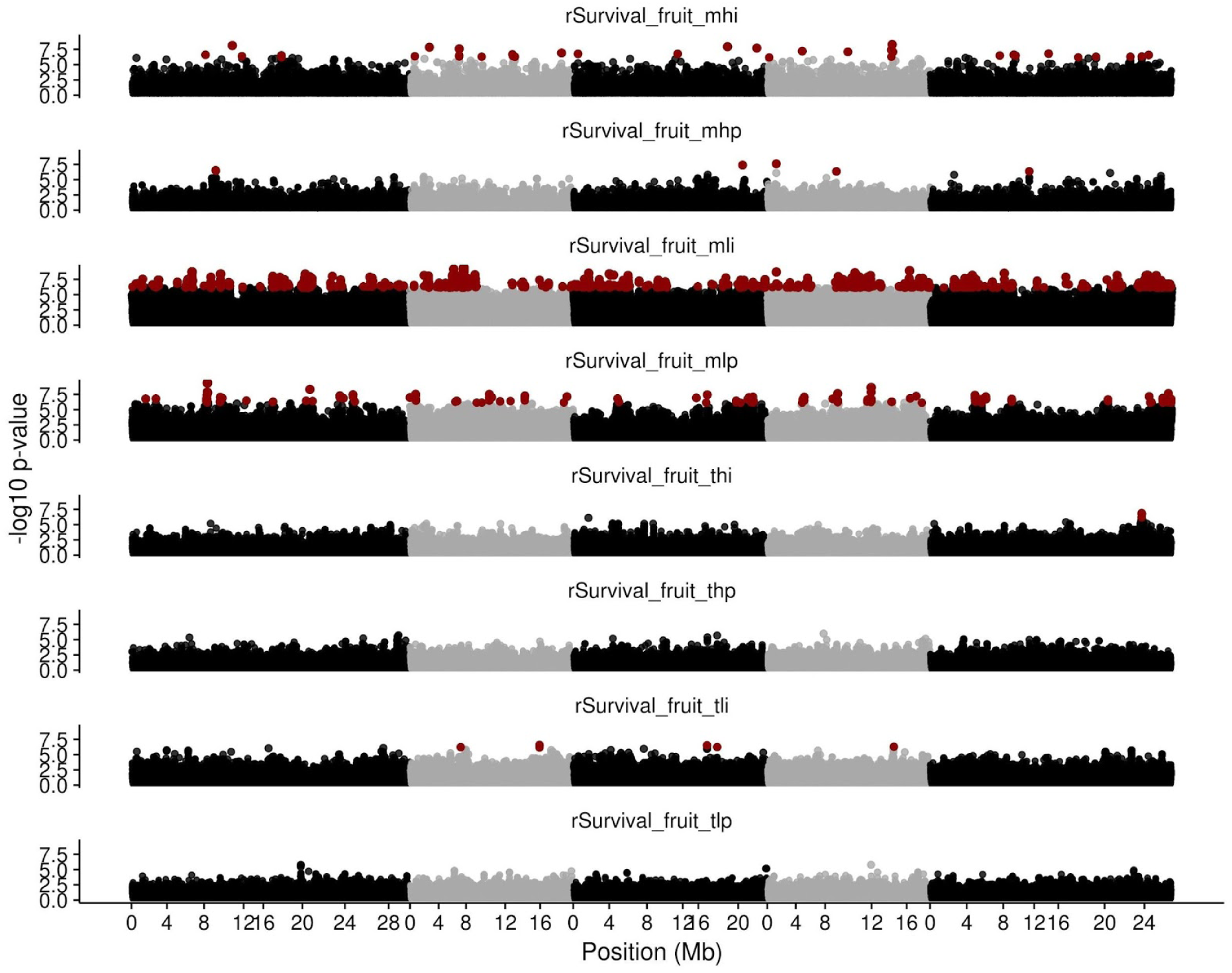
Genome maps of survival. Same as Fig. 1, but only using the survival component of fitness. [Abbreviations: The three characters of the codes: MLI, MLP, MHI, MHP, TLI, TLP, THI, TLP; indicate M=Madrid (Spain), T=Tübingen (Germany), L=Low precipitation, H=High precipitation, I=Individual replicates (one plant per pot), P=Population replicates (up to 30 plants per pot)].

**Figure SI.5.**
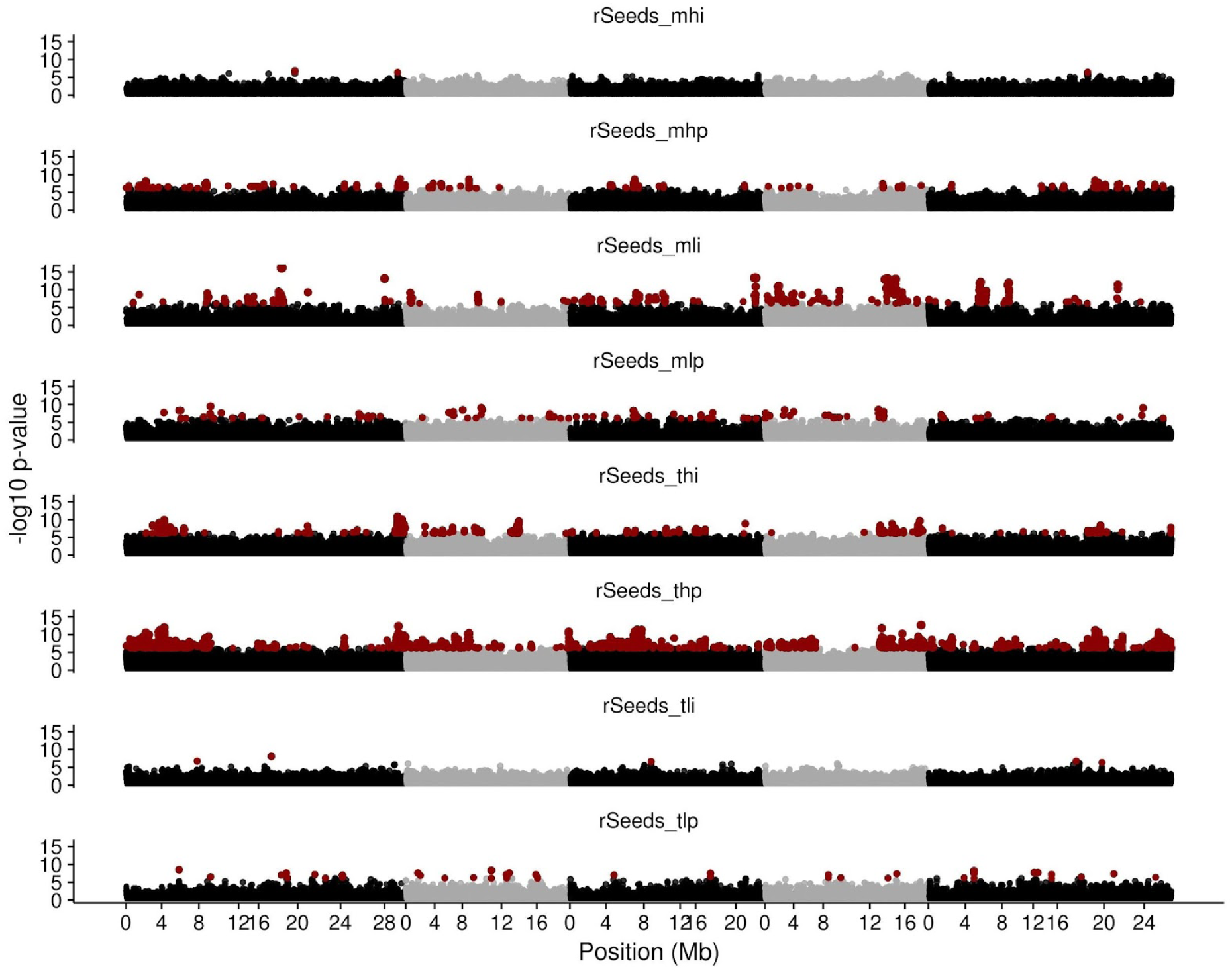
Genome maps of fecundity. Same as Fig. 1, but only using the fecundity component of fitness. [Abbreviations: The three characters of the codes: MLI, MLP, MHI, MHP, TLI, TLP, THI, TLP; indicate M=Madrid (Spain), T=Tübingen (Germany), L=Low precipitation, H=High precipitation, I=Individual replicates (one plant per pot), P=Population replicates (up to 30 plants per pot)].

**Figure SI.6.**
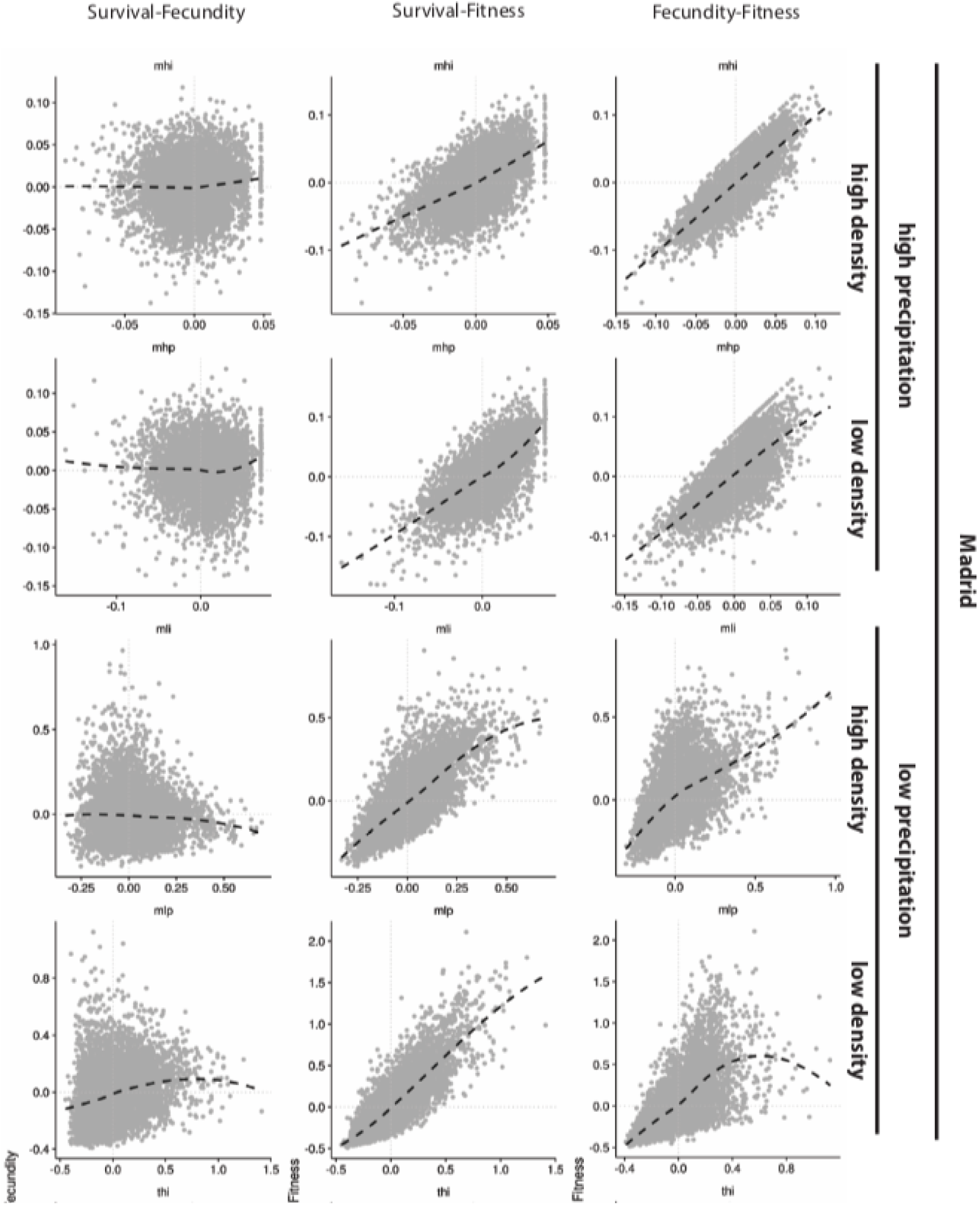
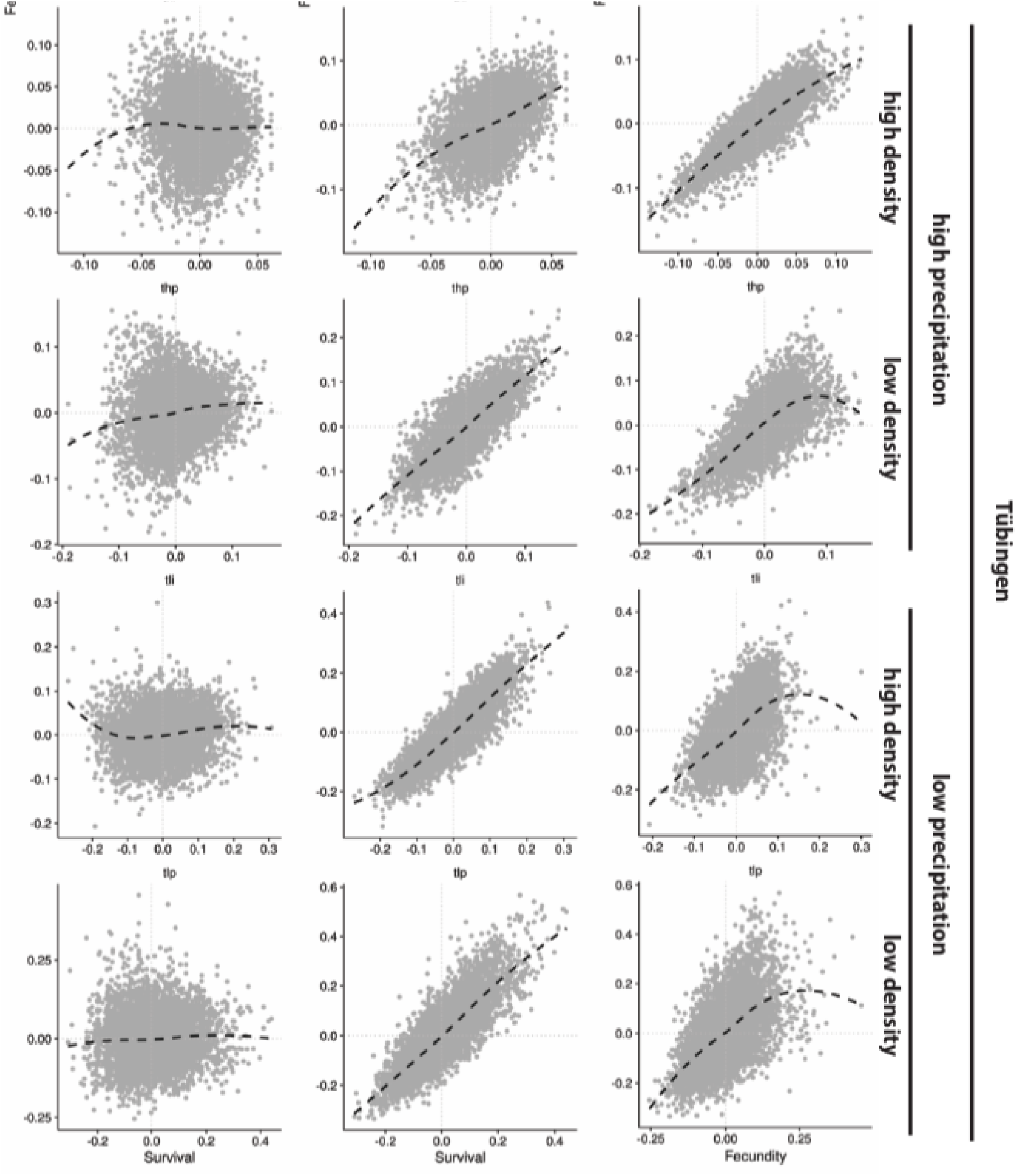
Trade-offs in survival and fecundity. Comparisons of total selection coefficients computed only with the survival component, only with the fecundity component, and with lifetime fitness. All environment combinations are plotted: Madrid (Spain) and Tübingen (Germany), high and low precipitation treatments, and high and low plant density treatments.

**Figure SI.7.**
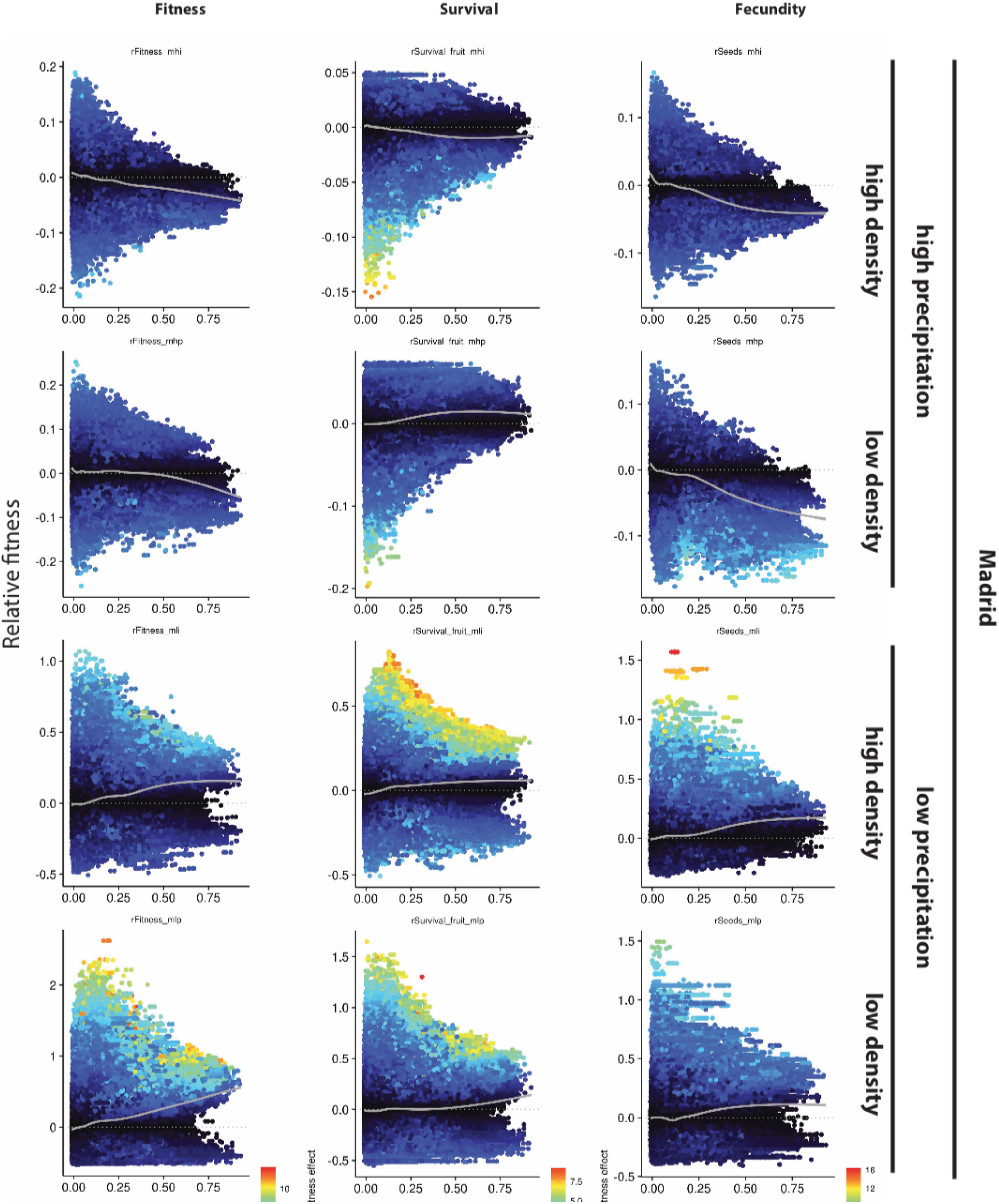
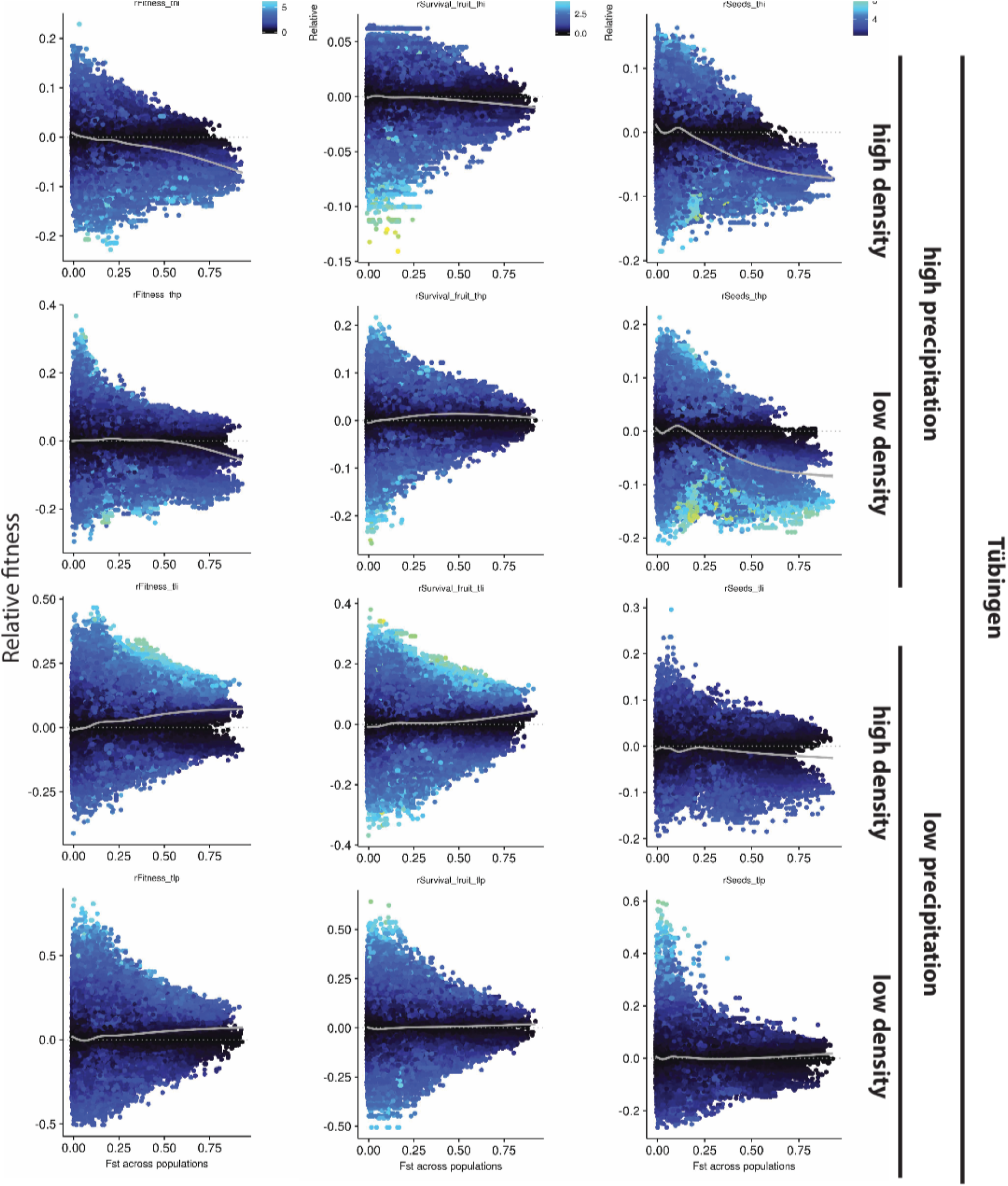
F_st_ and empirical selection. As Fig. 2C, for all environments: Madrid (Spain) and Tübingen (Germany), high and low precipitation treatments, and high and low plant density treatments.

**Figure SI.8.**
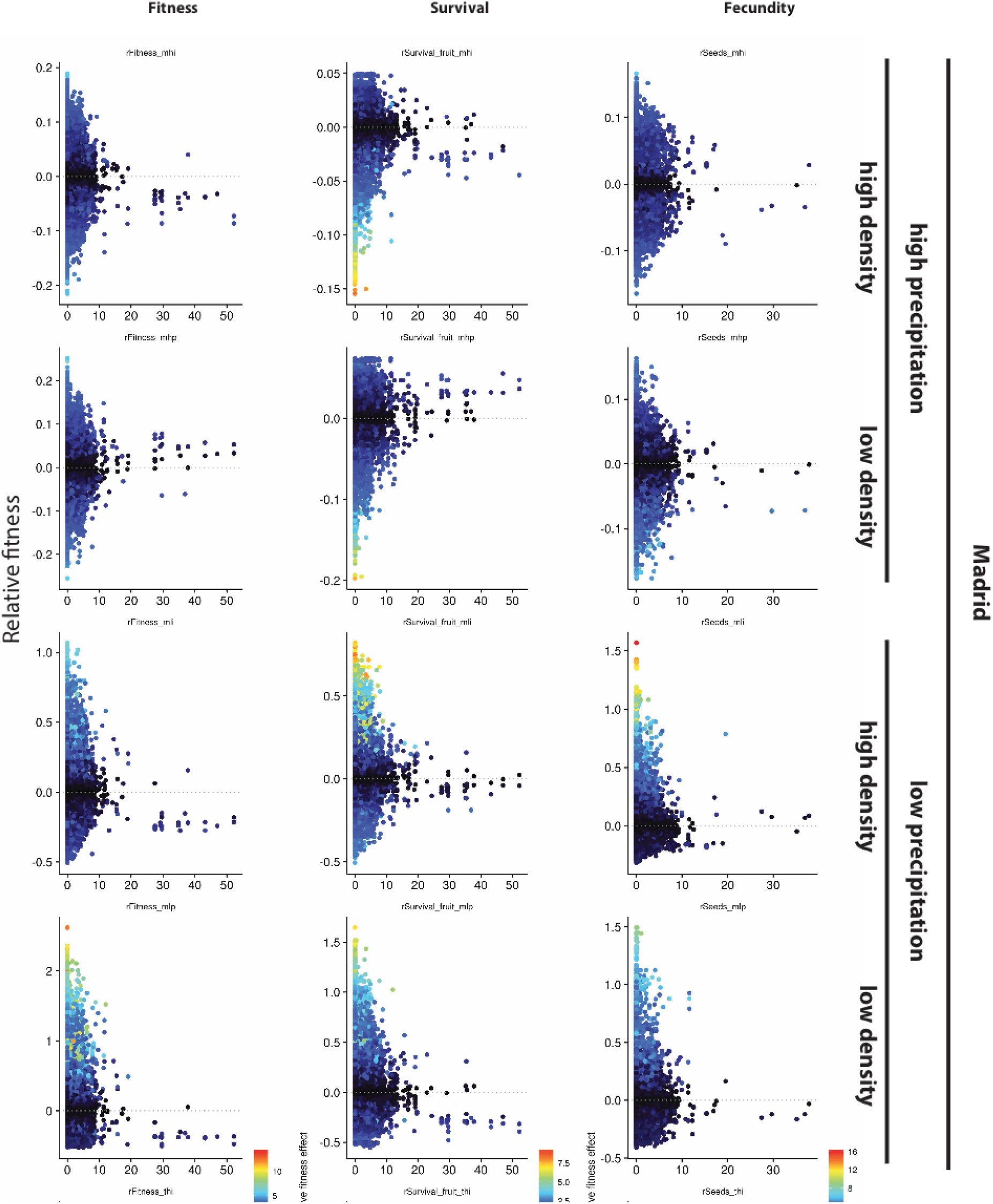
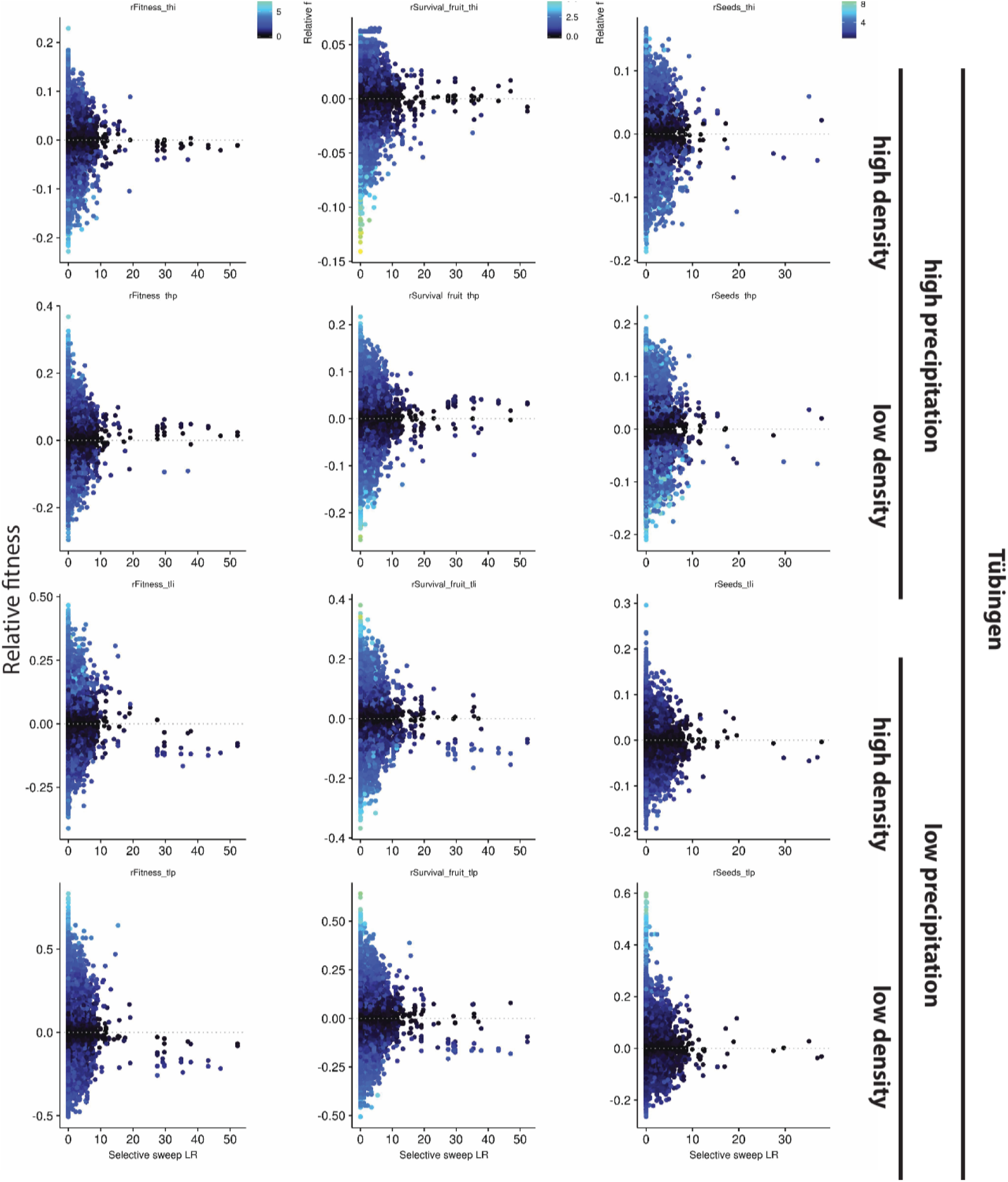
Sweeps and empirical selection. As Fig. 2D, for all environments: Madrid (Spain) and Tübingen (Germany), high and low precipitation treatments, and high and low plant density treatments.

**Figure SI.9.**
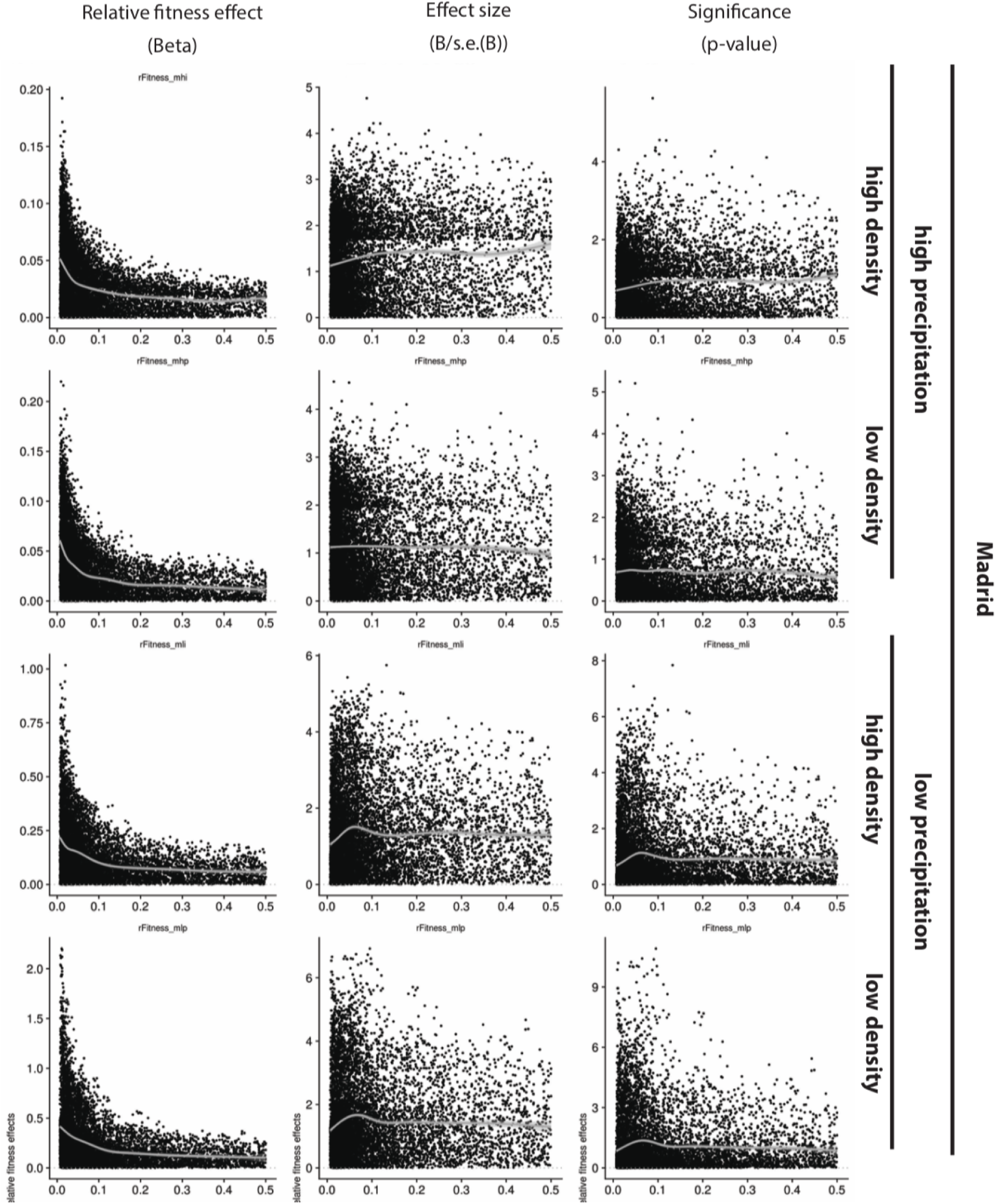
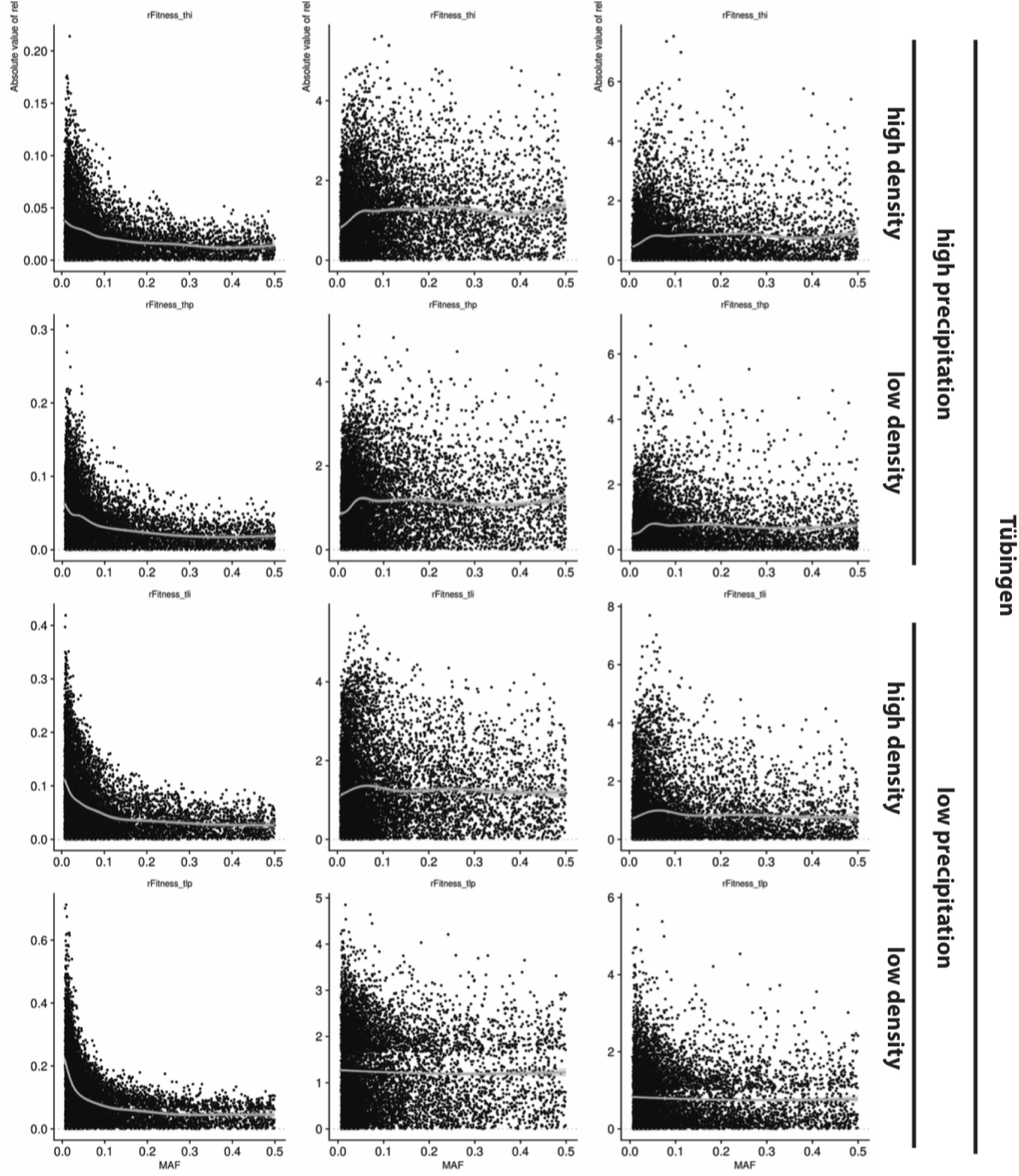
Allele frequency and empirical selection. Relationships between relative fitness effect, relative fitness effect size, and *P*-values (calculated from GWA with relative fitness) and minor allele frequency of alleles for all environments: Madrid (Spain) and Tübingen (Germany), high and low precipitation treatments, and high and low plant density treatments.

**Figure SI.10.**
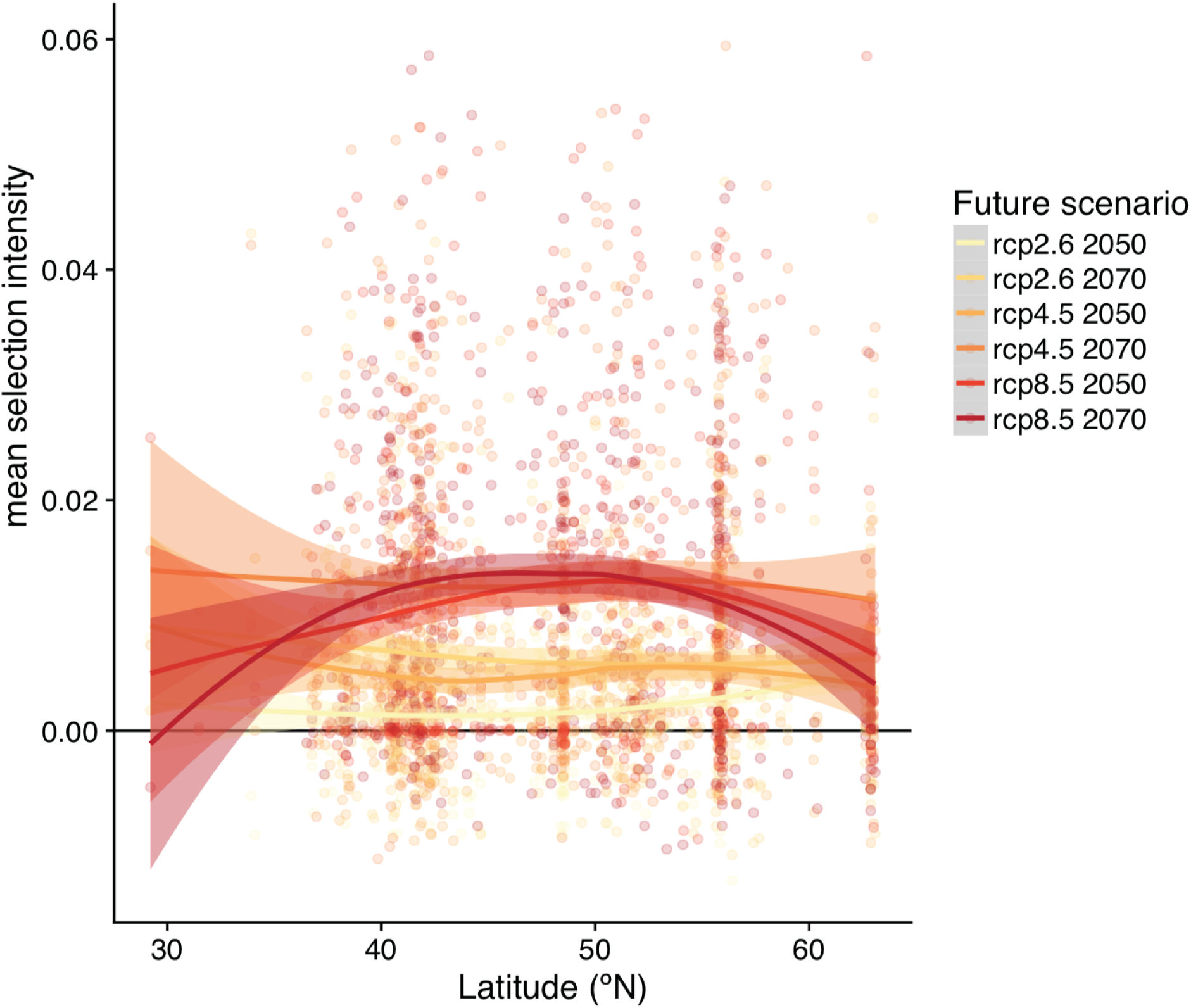
Future change in selection for different climate change scenarios. Same as Fig. 3G, but for different climate change scenarios. The higher the predicted CO_2_ emissions (rcp, representative concentration pathway), the stronger the predicted increase in selection intensity.

**Figure SI.11.**
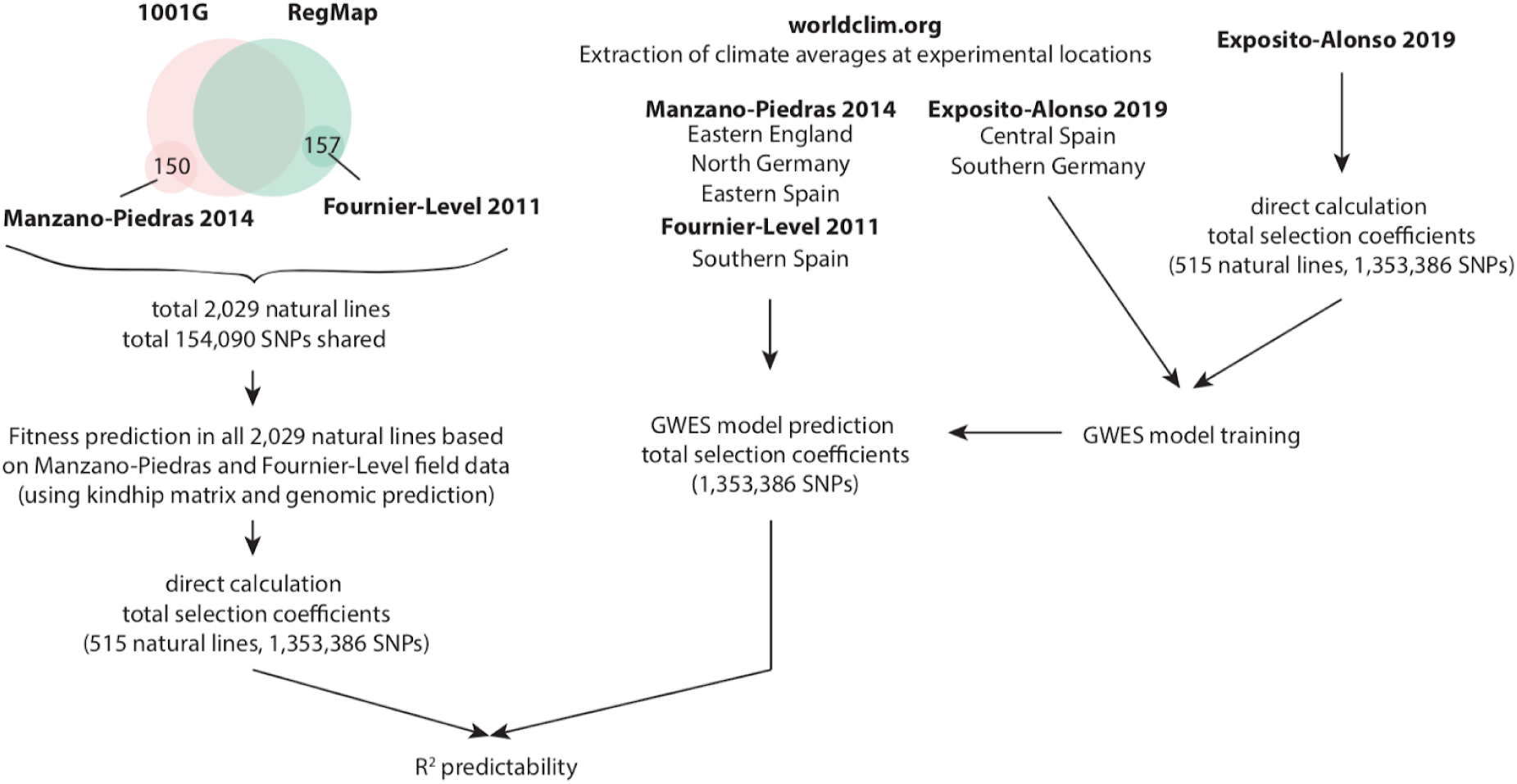
Field validation conceptual chart. Conceptual workflow on field validation procedure with data from published experiments (section VIII).

**Figure SI.12.**
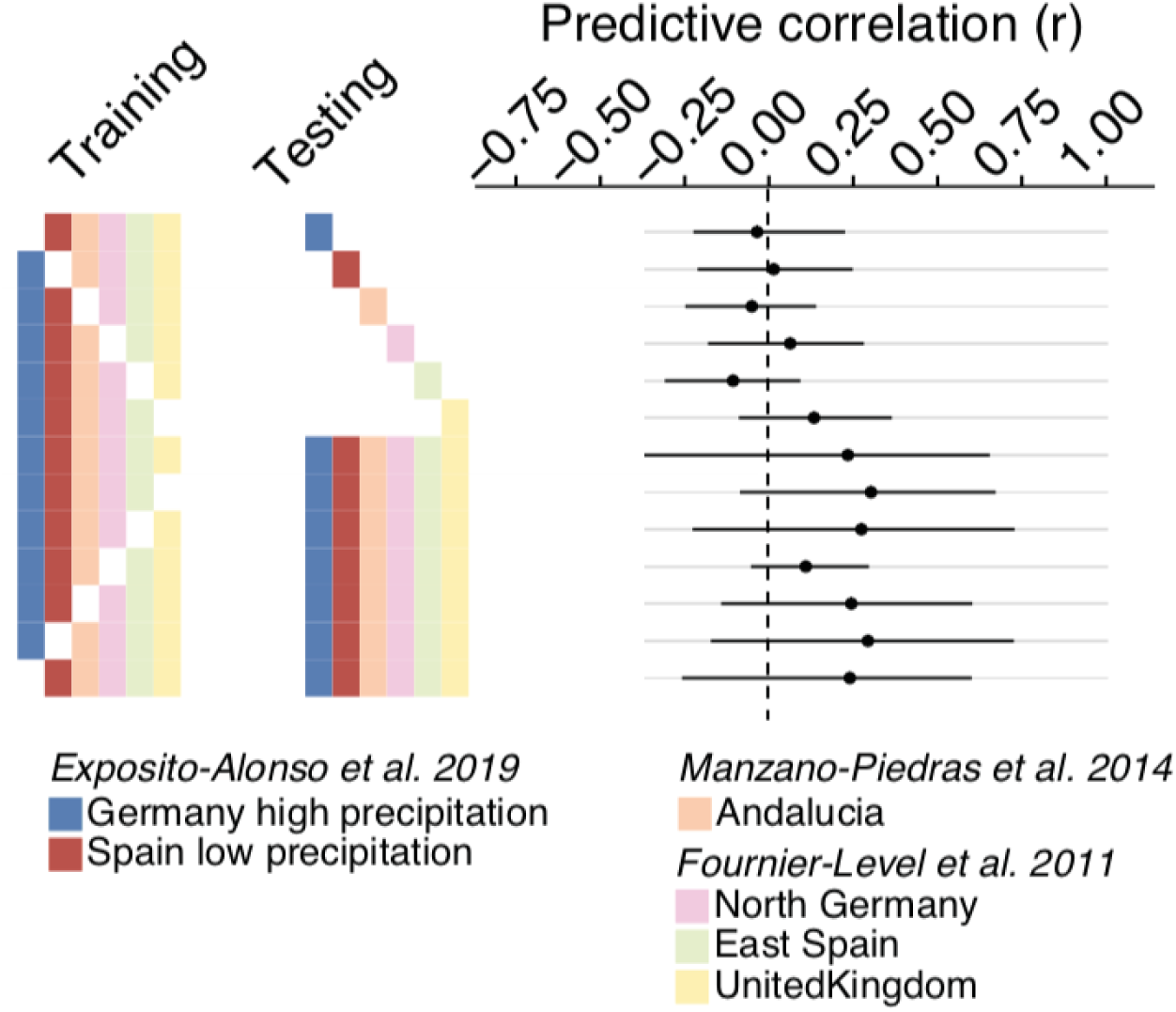
Null expectation of predictability. Same as Fig. 3, but with randomized fitness values associated to genotypes (section VIII). We could not find any model combination that had non-zero predictability (95% bootstrap confidence overlaps with zero). This proof of concept indicates that the predictability we find must have a biological basis, in which the combination of climate of origin for a genetic variant and the local climate allows to infer selection over such a variant.

**Figure SI.13.**
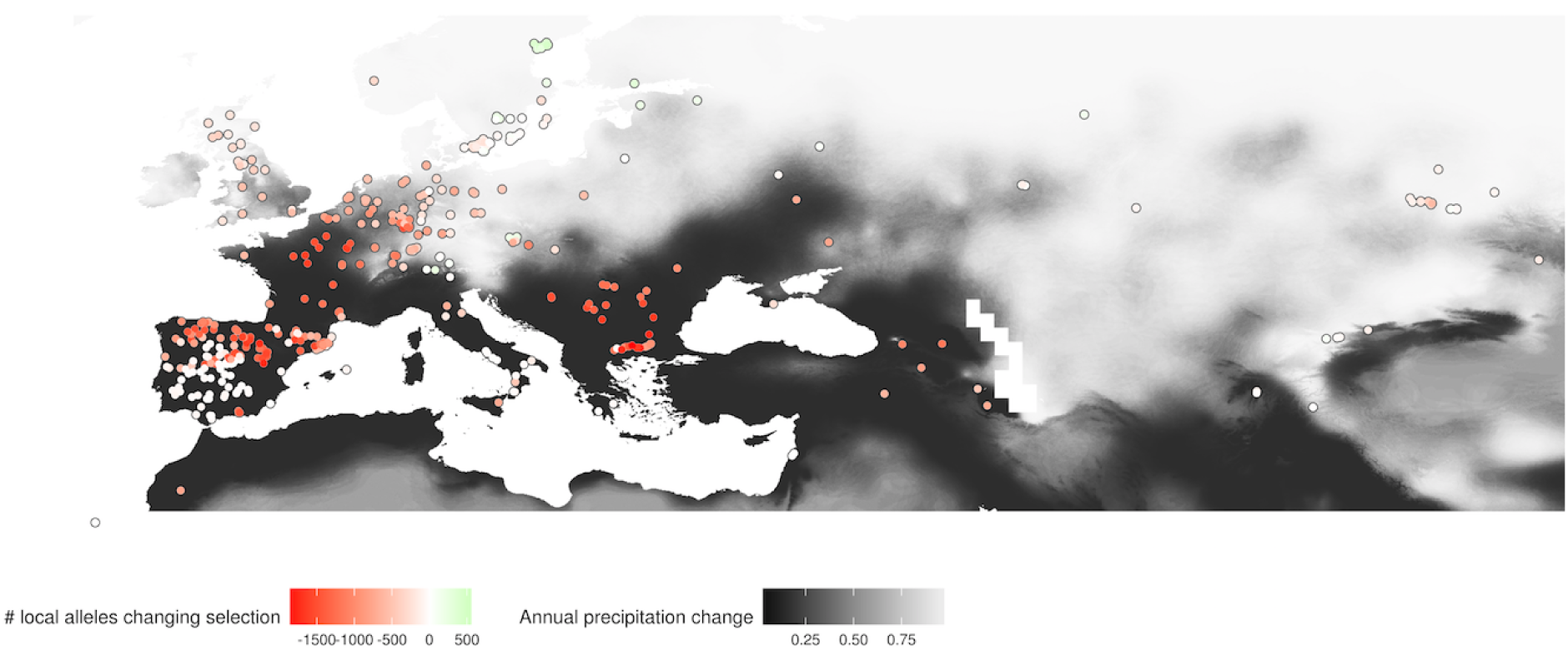
Change in selection relative to local diversity. Same as Fig. 3D, but counting the number of local alleles increasing or decreasing in selection (total n=10,752 SNPs). Only changes with more than 5% advantage/disadvantage were considered (defined *a posteriori* from Bonferroni-significant alleles, which generated at least 5% effect in fitness).

**Figure SI.14.**
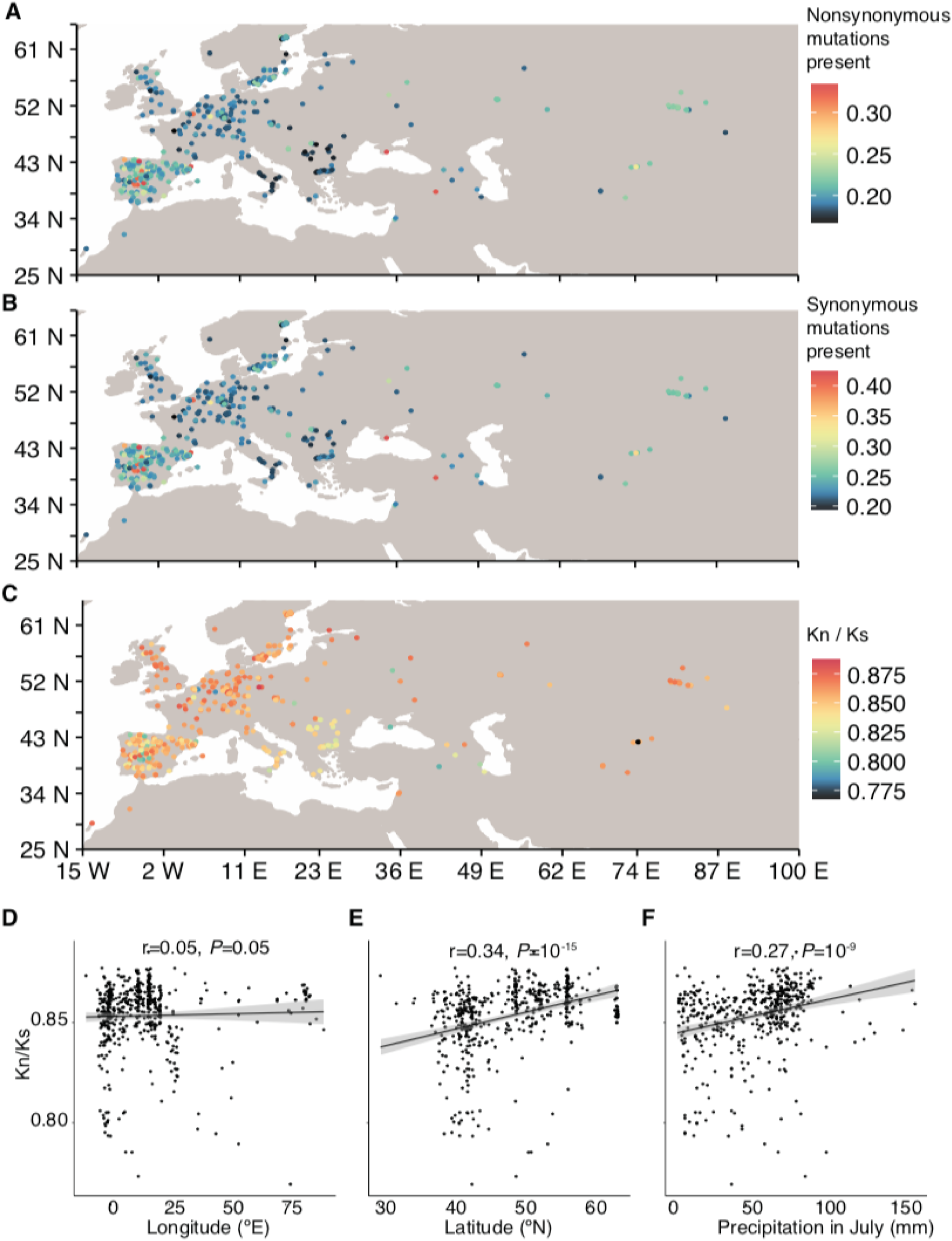
Deleterious and neutral mutations across space. Fraction of all genome-wide nonsynonymous (A) and synonymous (B) mutations present in the local genotype. (C) Ratio of nonsynonymous and synonymous fraction, i.e. K_n_K_s_. Correlation of K_n_K_s_ with degrees longitude, latitude, and precipitation in July (associated to selection intensity in Fig. 3). High selection intensity coincided with locations where natural lines have a lower-than-average ratio of nonsynonymous to synonymous polymorphisms (K_n_/K_s_, r=-0.276, 3×10^−10^), high local genetic diversity π (rho=0.187, *P*=2.63×10^−5^) and elevated Tajima’s D (rho=0.161, *P*=3×10^−4^). Various demographic scenarios could partially explain some of these patterns in isolation, i.e. bottlenecks can reduce the nonsynonymous polymorphisms because they are typically at low frequency, or high diversity might be found in old, large populations. These patterns are also in agreement with stronger selection having acted more efficiently over nonsynonymous mutations. In addition, high diversity could be driven by strong natural selection fluctuating over time, with alternative polymorphisms having been selected in each period. All in all, we did not find evidence that the warm edge of the geographic distribution of A. *thaliana* is formed by an increase in drift, which would cause small populations to accumulate nonsynonymous deleterious mutations and become genetically depauperate, as some theories propose^52^. Rather, our observations and predictions (Fig. 3) indicate that the species’ warm geographic limit is primarily defined by the environmental tolerance limits, where climate-driven natural selection limits the survival of individuals and populations outside the species range and only a few, highly specialized genotypes can survive near the range edges^53^.

**Figure SI.15.**
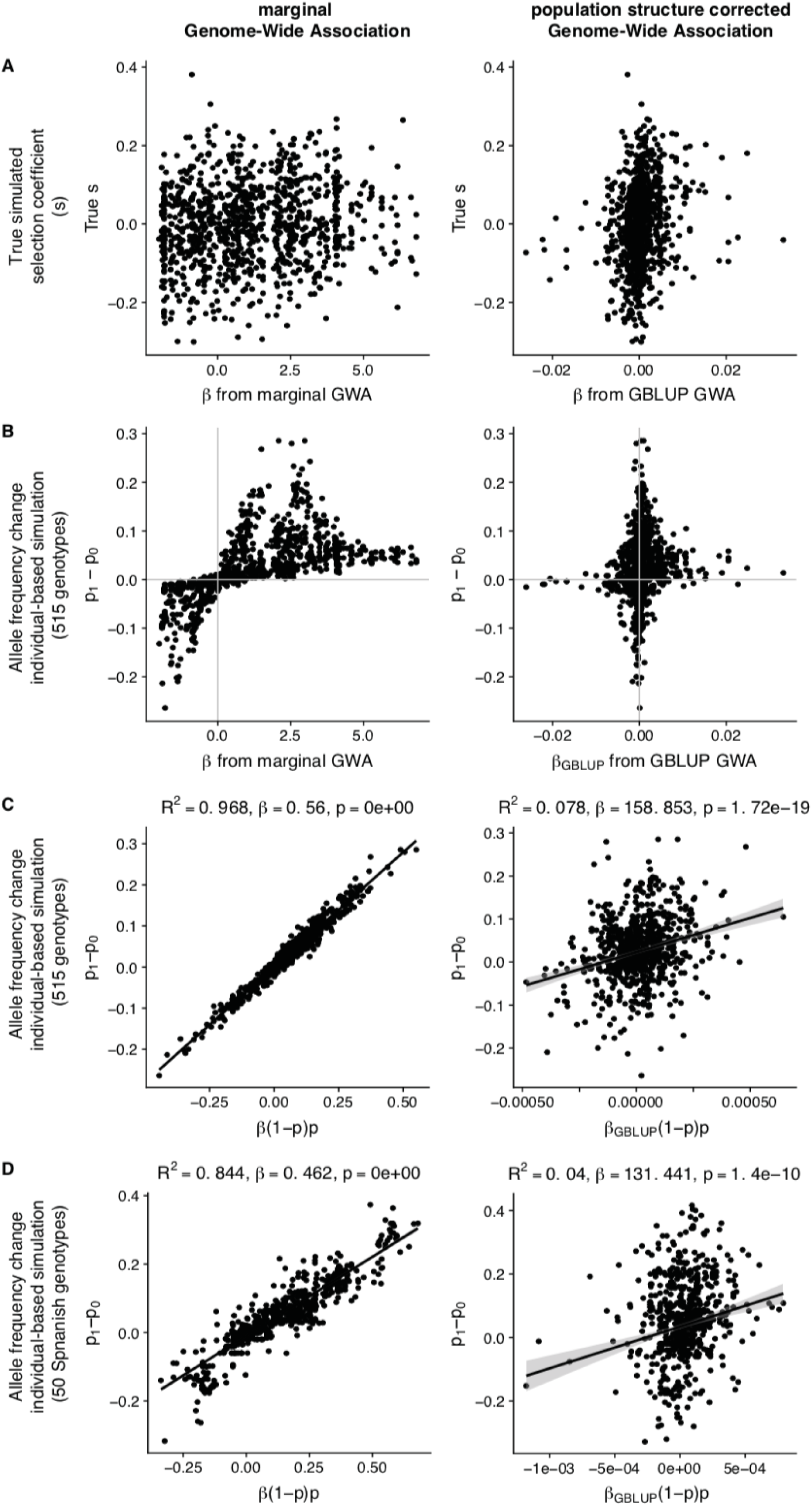
GWA model comparison in a simulation study of selection. We simulated fitness of 515 plants that differ in 1000 SNPs (subset of the original 1,353,386 genome matrix used throughout the manuscript) with selection coefficients drawn from a normal distribution around zero (more details and code are available at https://github.com/MoisesExpositoAlonso/selectioncorrelatedgenotvpes/, with doi: https://doi.org/10.5281/zenodo.1408095). (A) Comparison of simulated (true) values of selection coefficients and estimates from marginal GWA and GBLUP-based GWA. In the main text, these are called “total selection coefficients” and “direct selection coefficients", respectively. (B) Genotypes were sampled based on their relative fitness values to produce a population one generation after selection. Genome-wide allele frequency changes from generation zero (p_0_) to one generation after selection (p_1_) are compared to marginal and GBLUP GWA estimates. (C) We plug in GWA estimates into the theoretical equation of allele frequency change based on selection coefficients, Δ*p*= *P*(1 − *P*)^*s*^, and compare the theoretical and the simulated allele frequency changes in one generation. (D) In order to demonstrate extrapolability, we repeated (C) but instead of running the one-generation simulation of allele frequency with the 515 genotypes, we do so with only 50 Spanish genotypes. We repeat again the comparison of theoretical frequency changes based on GWA estimates with 515 genotypes, with the simulated allele frequency changes with 50 genotypes. All in all, the comparisons above indicate that marginal GWA estimates are appropriate to understand the consequences of selection in changing allele frequencies, even when extrapolating to other populations with slightly different allele frequencies and linkage patterns.

## SUPPLEMENTAL TABLES I

Supplemental tables are available in the online version of the paper with doi [update for publication] and are also deposited at Figshare with doi: https://doi.org/10.6084/m9.figshare.6756836.

**Table SI. 1. Summary of fitness data**

Average survival, fecundity, and lifetime fitness. Total number of genotypes with at least one surviving replicate per experiment.

[Abbreviations: The three characters of the codes: MLI, MLP, MHI, MHP, TLI, TLP, THI, TLP; indicate M=Madrid (Spain), T=Tübingen (Germany), L=Low precipitation, H=High precipitation, I=Individual replicates (one plant per pot), P=Population replicates (up to 30 plants per pot)].

**Table SI.2. Heritability of fitness**

Broad sense heritability and the 95% Highest Posterior Density Interval per trait (variance explained by line genotype), as calculated from a generalized linear mixed model using MCMCglmm, is reported as: *σ*_*g*_/*σ*_*Total*_. The proportion of variance explained by nuisance factors such as block (tray), position of the tray within a treatment block, and position of plant within a tray are reported in the same way. Proportion of Variance Explained (chip heritability) and the 95% Highest Posterior Density Interval per trait, as calculated from a Bayesian Sparse Linear Mixed Model (BSLMM-GEMMA) using genotype means per trait and using a kinship/relationship matrix.

[Abbreviations: The three characters of the codes: MLI, MLP, MHI, MHP, TLI, TLP, THI, TLP; indicate M=Madrid (Spain), T=Tübingen (Germany), L=Low precipitation, H=High precipitation, I=Individual replicates (one plant per pot), P=Population replicates (up to 30 plants per pot)].

**Table SI.3. Number of SNPs with significant total selection coefficients**

All significant variants from marginal GWA after FDR and Bonferroni correction and all variants with non-zero probability of inclusion from conditional GWA, and sharing of significant variants across experiments.

**Table SI.4. Expected allele frequency changes in response to selection**

Summaries of allele frequency changes per experiment.

**Table SI.5. Odds ratio of antagonistic pleiotropy vs conditional neutrality**

Fisher’s Exact Test Odds Ratio (OR) and *P*-values of two complementary tests: pleiotropy vs conditional neutrality (CNAP column) and antagonistic vs non-antagonistic pleiotropy (APnAP). This was performed for combined lifetime fitness, survival, and fecundity.

**Table SI.6. Correlation of total selection coefficients across environments**

**Table SI.7. Variable importance of predictive models**

**Table SI.8. Predictability of environmental models**

After training GWES models with a set of experiments, we inferred total selection coefficients on another set of experiments and compared those with the real total selection coefficients. We calculated Pearson’s product-moment correlation r_cv_ and percentage of variance explained R^2^ _cv_ using a regression. 95% confidence intervals were calculated with 100 bootstrap replicates.

[Abbreviations: ml= Central Spain and low precipitation (both high and low plant density treatments combined); th= South Germany and high precipitation (both high and low plant density treatments combined), Andalucia= South Spain from Manzano-Piedras *et al.* (2014, ref. ^26^), Germany= North Germany from Fournier-Level *et al.* (2011, ref. ^25^), Spain= South East Spain from from Fournier-Level *et al.* (2011, ref. ^25^), United Kingdom= East England from Fournier-Level *et al.* (2011, ref. ^25^)].

**Table SI.9. Description of climate variables**

Climate variables used for environmental models are described and their sources reported.

**Table SI.10. GBLUP heritability and imputation accuracy of published field data**

We used GBLUP to impute fitness from Fournier-Level *et al.* (2011) and Manzano-Piedras *et al.* (2014) into our 517 global accessions. We report heritability, Pearson’s r between GBLUP predicted fitness and real fitness, and the significance of the correlation test.

**Table SI.11. Correlation between inferred natural selection intensity and other variables**

Spearman’s rho between selection intensity and diversity metrics or climate metrics is given.

## SUPPLEMENTAL APPENDIX II: A rainfall-manipulation experiment with 517 *Arabidopsis thaliana* accessions

Moises Exposito-Alonso^1^, Rocío Gómez Rodríguez^2^, Cristina Barragán^1^, Giovanna Capovilla^1^, Eunyoung Chae^1^, Jane Devos^1^, Ezgi S. Dogan^1^, Claudia Friedemann^1^, Caspar Gross^1^, Patricia Lang^1^, Derek Lundberg^1^, Vera Middendorf^1^, Jorge Kageyama^1^, Talia Karasov^1^, Sonja Kersten^1^, Sebastian Petersen^1^, Leily Rabbani^1^, Julian Regalado^1^, Lukas Reinelt^1^, Beth Rowan^1^, Danelle K. Seymour^1^, Efthymia Symeonidi^1^, Rebecca Schwab^1^, Diep Thi Ngoc Tran^1^, Kavita Venkataramani^1^, Anna.Lena Van de Weyer^1^, François Vasseur^1^, George Wang^1^, Ronja Wedegärtner^1^, Frank Weiss^1^, Rui Wu^1^, Wanyan Xi^1^, Maricris Zaidem^1^, Wangsheng Zhu^1^, Fernando García-Arenal^2^, Hernan A. Burbano^1^, Oliver Bossdorf^3^, Detlef Weigel^1^.

^1^Department of Molecular Biology, Max Planck Institute for Developmental Biology, Tübingen, Germany. ^2^Center for Plant Biotechnology and Genomics, Technical University of Madrid, Pozuelo de Alarcón, Spain. ^3^Institute of Ecology and Evolution, University of Tübingen, Tübingen, Germany.

### I. Background & Summary

The gold standard for studying natural selection and adaptation in the wild is to quantify lifetime fitness of individuals from natural populations that have been grown together in a common garden, or that have been reciprocally transplanted. Natural selection over morphological, physiological or other traits has been studied in a wide range of organisms^13,54-56^ using observational and experimental fitness measurements of multiple individuals in field conditions. However, studies that combine such measurements with knowledge on genome-wide variation are, in comparison, very rare^3,15,22^. This is surprising, given that they would enable the translation of selection to the genetic level and thus ultimately help us to understand whether traits will evolve over generations.

With climate change, the study of adaptation to the environment has acquired new importance. Predictions of climate change indicate not only that temperature will rise, but that also precipitation regimes will be altered, leading to more frequent and extreme droughts^57^ and seriously threaten the persistence of plant communities^2,58^. Field experiments where climate variables such as rainfall are manipulated can be used to address this question^59^.

Here we present a high-throughput field experiment with 517 whole-genome sequenced natural lines of *Arabidopsis thaliana^16^.* This experiment was designed to be of a sufficiently large scale to enable powerful genome-wide association analyses^60^ and to maximize the replicability of species-wide patterns, which has been shown to increase with the diversity of genotypes included in an experiment^61^. The experiments were conducted in two field stations with contrasting climate, in the Mediterranean (Spain) and in Central Europe (Germany), where we built rainout shelters and simulated high and low rainfall. Using custom image analysis we quantified fitness- and phenology-related traits for 23,154 pots, which contained about 14,500 plants growing independently, and over 310,000 plants growing in small populations (max. 30 plants per pot). Three measurements of fitness were produced: survival from seed to reproductive adult (proportion 0—1) and the average fecundity per reproductive adult (inflorescence skeleton lengths ranged from 18,400 to 1,622,000 pixels, which approximately corresponds to 1 to 6,127 seeds per plant). Fecundity was only measured for plants with at least one fruit. We finally calculated an integrated lifetime fitness value by multiplying the survival proportion to adulthood with the total offspring produced. This dataset will be invaluable for the study of natural selection and adaptation in the context of global climate change at the genetic level, building on the genetic catalog of the 1001 Genomes Project^16^ and complementing the already published extensive set of traits measured in controlled growth chamber or greenhouse conditions^62,63^.

### II. Selection of accessions from the 1001 Genomes Project

The 1001 Genomes (1001G) Project^16^ has provided information on 1,135 natural lines or accessions and 11,769,920 SNPs and small indels called after re-sequencing. To select the most genetically and geographically informative 1001G lines, we applied several filters: (1) First we removed the accessions with the lowest genome quality. We discarded those with < 10X genome coverage of Illumina sequencing reads and < 90% congruence of SNPs called from MPI and GMI pipelines^16^. (2) We removed near-identical individuals. Using PLINK software^33^ we computed identity by state across the 1,135 accessions. For pairs of accessions with < 0.01 differences per SNP (<100,000 variants approx.), we randomly selected one accession to include in our study. (3) Finally, we reduced geographic sampling ascertainment bias, as the sampling for 1001G was performed in neither a random nor a regularly structured scheme. Some laboratories provided several lines per location whereas others provided lines that were collected at least several hundred kilometers apart. Using each accession’s collection location, we computed Euclidean distances across the 1,135 accessions and identified all pairs that were apart less than 0.0001 Euclidean distance in degrees latitude and longitude (<< 100 meters). From such pairs, we randomly selected one accession to remain. After applying criteria (1), (2), and (3), we obtained a final set of 523 accessions (<u>Datasets 1 and 2</u>). To bulk seeds for our rainfall-manipulation experiment and control for maternal effects, we first propagated accessions in controlled conditions. We stratified the seeds one week at 4°C, we sowed them in trays with industrial soil (CL-P, Einheitserde Werkverband e. V., Sinntal-Altengronau Germany) and placed them in a growth room with 16 h light and 23°C for one week. Trays were vernalized for 60 days at 4°C and 8 h day length. After vernalization, trays were moved back to 16 h light and 23°C for final growth and reproduction. This generated sufficient seeds for 517 accessions, which were later grown in the field in two locations (Fig. SII.1). Seeds originating from the same parents can be ordered from the 1001G seed stock at the Arabidopsis Biological Resource Center (CS78942).

### III. Field experiment design

#### III.1 Rainout shelter, watering, and block design

We built two 30 m × 6 m tunnels of PVC plastic foil to fully exclude rainfall in Madrid (Spain, <u>40.40805ºN −3.83535ºE</u>) and in Tübingen (Germany, <u>48.545809ºN 9.042449ºE</u>) (Fig. SII.2A-B). The foil tunnels are different from a regular greenhouse in that they are completely open on two sides. Thus, ambient temperatures vary virtually as much as outside the foil shelter (see Environmental sensors section). In each location, we supplied artificial watering in two contrasting regimes: abundant watering and reduced watering. Inside each tunnel, we created a 4% slope, and four flooding tables (two for high and two for low precipitation) (1 m × 25 m, Hellmuth Bahrs GmbH & Co KG, Brüggen, Germany) covered with soaking mats (4 l/m^2^, Gärtnereinkauf Münchingen GmbH, Münchingen, Germany). The flooding tables were placed on the ground in parallel to the slope. Water was able to drain at the lower end of the flooding table (Fig. SII.2A-B). A watering gun was used to manually simulate rainfall from the top.

Our experimental design is a split-plot design (Fig. SII.2C), with precipitation treatments replicated twice in each location and the genotypes randomized within precipitation treatment in a total of 8 spatial blocks. This ensured that all genotypes would be equally evenly distributed within the foil tunnel, and that we could robustly measure consistent fitness responses to water deprivation across precipitation replicates.

On top of the flooding tables, we used potting trays with 8×5 cells (5.5 cm × 5.5 cm × 10 cm size) and industrial soil (CL-P, Einheitserde Werkverband e.V., Sinntal-Altengronau Germany). Each cell would correspond to a genotype, excluding corner cells, to avoid extreme edge effects. We grew a total of 12 replicates per genotype per treatment: Five replicates were grown at high density, with 30 seeds per cell and without further intervention ("population replicate"). The remaining seven replicates were at low density (ca. 10 seeds) and one seedling was selected at random after germination ("individual replicate"). Excess individuals were culled. While the population replicates should more faithfully reflect survival from seed to reproduction, the individual replicates were useful to more accurately monitor flowering time and seed set.

#### III.2 Environmental sensors

Environmental variables — air temperature, photosynthetically active radiation (PAR) and soil water content — were monitored every 15 minutes for the entire duration of the experiment using multi-purpose sensors (Flower Power, Parrot SA, Paris, France). This enabled us to adjust watering depending on the degree of local evapotranspiration during the course the experiment. The sensors outside of the tunnel in Madrid (i.e. only natural rainfall) showed an interquartile range between 1% and 17% soil water content. This overlapped with the range of 10 to 22% water content of the drought treatment that we artificially imposed inside the tunnels in Madrid and Tübingen. The lower range of measurements in Madrid (outside sensor) is due to a lack of natural rainfall during the first two months of the experiment (Fig. SII.2E, Table SII.1). In contrast, the sensor outside the tunnel in Tübingen recorded an interquartile range of soil water content percentage of 22 to 27%, which was comparable to the high watering treatments in Tübingen and Madrid (from 20 to 33%) (Fig. SII.2E, Table SII.1). These values confirmed that our low and high watering treatment were not only different, but also that they mimicked natural soil water content at the two contrasting locations. Mean daily air temperatures (measured by the sensors at 5-10 cm above the soil surface every 15 minutes) were overall higher in Madrid (8-10°C) than in Tübingen (5-6°C), and the difference in temperature between the sensors inside and outside the tunnels was in both locations on average only 1°C (Fig. SII.2F, Table SII.1). The photosynthetically active radiation (PAR, wavelengths from 400 to 700 nm) had a median of 0.1 mol m^−2^ day^−1^ at night for all experiments. At mid-day (11:00-13.00 hrs), the median PAR in Madrid was 57.8 mol m^−2^ day^−1^ outside, and 45.7 mol m^−2^ day^−1^ inside the tunnel. In Tübingen, the median values were 29.0 outside, and 30.9 mol m^−2^ day^−1^ inside the tunnel.

#### III.3 Sowing and quality control

During sowing, contamination of neighbouring pots with adjacent genotypes can occur for multiple reasons. In order to avoid such contamination, we chose a day with no wind and sowed seeds at 1-2 cm height from the soil. Additionally, we took care during the first days to be particularly gentle when using the watering gun to avoid seed-carryover (bottom watering by flooding was done regularly). We also tried to remove human error during sowing by preparing and randomizing 2 ml plastic tubes containing the seeds to be sown in the same layouts (5×8) as the destination trays. During sowing, each experimenter took a box at random and went to the corresponding labeled and arranged tray in the field (Fig. SII.2). This reduced the possibility of sowing errors. Sowing occurred on November 16 2015 in Madrid and on October 22 2015 in Tübingen. During vegetative growth, we could identify seedlings that resembled their neighbours or were located in the border between two pots and removed such plants as potential contaminants. We also used the homogeneity of flowering within a pot in the population replicates as a further indicator for contamination (Fig. SII.3A). When a plant had a completely different flowering timing or vegetative phenotypes did not coincide with the majority of plants in the pot, this plant was removed. After sowing and quality control, the total number of pots was 24,747 instead of the original 24,816 pots (99.7%) (<u>Dataset 3</u>).

### IV. Field monitoring

#### IV.1 Image analysis of vegetative rosettes

Top-view images were acquired every four to five days (median in both sites) with a Panasonic DMC-TZ61 digital camera and a customized closed dark box, the “Fotomatón” (Fig. SII.3A), at a distance of 40 cm from each tray. In total, we imaged each tray at 20 timepoints throughout vegetative growth. The implemented segmentation was the same as in Exposito-Alonso *et al.^8^,* which relies on the Open CV Python library^64^. We began by transforming images from RGB to HSV channels. We applied a hard segmentation threshold of HSV values as (H=30-65, S=65-255, V=20-220). The threshold was defined after manually screening 10 different plants in order to capture the full spectrum of greens both of different accessions and of different developmental stages. This was followed by several iterations of morphology transformations based on erosion and dilation. For each of the resulting binary images we counted the number of green pixels.

During field monitoring, we noticed that some pots were empty because seeds had not germinated. In these cases, we left a red marker in the corresponding pots (Fig. SII.3A),which could be detected in a similar way as the presence of green pixels (with threshold H=150-179, S=100-255, V=100-255). These pots were excluded from survival analysis as they did not contain any plants (Fig. SII.3A). The resulting raw data consist of green and red pixel counts per pot (Fig. SII.3B). In order to detect the red markers automatically, we performed an analysis of variance between pots above and below a threshold of red pixels and finding the threshold that maximized this separation (Fig. SII.3C). This provided us with the threshold of red pixels above which a pot had a red marker (indicating an empty pot). As expected, the distribution of pixels was bimodal, making this identification straightforward.

We estimated germination timing by analysing trajectories (Fig. SII.3B) of green pixels per pot, and identifying the first day that over 1,000 green pixels were observed in a pot (corresponding to a plant size of ~ 10 mm^2^, Fig. SII.3) (<u>Datasets 3</u>). The final dataset contained data for 22,779 pots — after the removal of pots with red labels — with a time series of green pixel counts.

#### IV.2 Manual recording of flowering time

We visited the experimental sites every 1-2 days and manually recorded the pots with flowering plants. Flowering time was measured as the day when the first white petals could be observed with the unaided eye. This criterion was chosen as sufficiently objective to reduce experimenter error. To keep track of previous visits and avoid errors, we labeled the pots where flowering had already been recorded with blue pins. To calculate flowering time, we counted the number of days from the date of sowing to the recorded flowering date (we did not use the inferred day of germination to avoid introducing modeling errors in the flowering time metric). Fig. SII.4A shows the raw flowering time data per pot in the original spatial distribution and the distribution of flowering time per treatment combination. Note that grey boxes are pots with plants that did not survive until flowering. In total, we gathered data for 16,858 pots with flowering plants (<u>Datasets 3</u>).

#### IV.3 Image analysis of reproductive plants

Once the first dry fruits were observed, we harvested them and took a final “studio photograph” of the rosette and the inflorescence (Fig. SII.5A). In total, we took 13,849 photographs. The camera settings were the same as for the vegetative monitoring, but here we included an 18% grey card approximately in the same location for each picture in case *a posteriori* white balance adjustments would be needed. We first used a cycle of morphological transformations of erode-and-dilate to produce the segmented image (Fig. SII.5C). This generated a segmented white/black image without white noise. Then, we used the thin (erode cycles) algorithm from the Mahotas Python library^65^ to generate a binary picture reduced to single-pixel paths — a process called skeletonisation (Fig. SII.5C). Finally, to detect the branching points in the skeletonised image we used a hit-or-miss algorithm. We used customized structural elements to maximize the branch and end point detection (Fig. SII.5C). This resulted in four variables per image: total segmented inflorescence area, total length of the skeleton path, number of branching points, and number of end points (Fig. SII.5C) (<u>Datasets 3-4</u>).

#### IV.4 Estimation of fruit and seed number

Although the study of natural selection is based on studying relative fitness, and total reproductive area might provide a good relative estimate, sometimes it is useful to have a proxy of the absolute fitness. In order to provide an approximate number of how many seeds each plant had produced, we generated two allometric relationships by visual counting of fruits per plant and seeds per fruit. In order to be sure that the counts corresponded to single plants, we counted fruits and seeds of only individual replicates of accessions, not the population replicates (see Field experiment design section). Because a strong relationship had already been validated between inflorescence size and the number of fruits in a number of studies with *A. thaliana^66–68^,* we decided that counting a few inflorescences of three sizes, reflecting the broad size spectrum, would be sufficient to establish a first allometric relationship with the four image-acquired variables (n=11 inflorescences, R^2^=0.97, P=4×10^−4^, Fig. SII.5B). To express fecundity as the number of seeds, we counted all seeds inside one fruit for each of the inflorescences used for the first allometric relationship (n=11 fruits), aiming for a wide range of fruit sizes. The mean was 28.3 seeds per fruit and the standard deviation was 11.2 seeds. The two aforementioned allometric relationships were used to predict, first, the number of fruits per inflorescence using the four image analysis variables, and second, the number of seeds corresponding to the number of fruits per inflorescence (Datasets 3-4).

### V. Technical validations

#### Data processing

All images, from where fruits and leaf area were estimated, are backed up and stored at the Max Planck Institute for Developmental Biology and available through ftp transfer (ca. 2Tb) upon request to. The Max Planck Society requires storage of publication-relevant data for a minimum of 10 years. The Python modules to process images for green area segmentation and inflorescence analyses are available at http://github.com/MoisesExpositoAlonso/hippo and http://github.com/MoisesExpositoAlonso/hitfruit, along with example datasets. The scripts for field data curation doi 10.5281/zenodo.2583224. The field data curation R package dryAR (http://github.com/MoisesExpositoAlonso/drvARl.

#### Replicability of image processing

After testing different camera parameters, we used an exposure of-2/3 and an ISO of 100. White balance was set for flashlight. We used a dark box with all sides closed, so the flashlight was the only source of illumination. This ensured that the white balance and illumination were virtually consistent from picture to picture, as shown before^8^. Photos were saved both in .jpeg and .raw to allow for *a posteriori* adjustments if needed. Using a calibration board with 1.3 cm × 1.3 cm white and dark squares, we examined the error between the inferred area from image analysis and the real 1.3 cm-side squares across the tray. This provided us with a median resolution estimate of 101.5 pixels mm^−2^. The deviations from the true area were minimal, with a median of 2.7% and values of 1.4% / 4.2% for the 1^st^ and 3^rd^ quartile. The maximum area deviations were of 8 to 9% in the extreme corners of the tray, where we did not sow any seeds. We are confident that such small variation in retrieved area is compensated by the randomized locations of genotypes within the trays.

To further verify that our camera settings and segmentation pipeline produced replicable extractions of plant green area, we used images of trays that were photographed twice on the same day by mistake. In total, there were 1,508 such pots distributed across 11 time points and different trays. By comparing the area of the same pot of two different camera shots and segmentation analyses, we could verify that the Spearman’s rho of rank correlation was very high (r=0.97, n=1508, *P*<10^−16^), confirming high replicability.

Because we ran the same segmentation and skeletonisation software on both rosette and inflorescence images, we could leverage the clearly different image patterns that rosettes and inflorescences have to identify labeling errors (i.e. mistakes in manually inputting sample information of the pictures). To do this, we first trained a random forest model to predict the manually labeled “rosette” or “inflorescence” by the four image variables (Fig.SII.5). By fitting a Random Forest with all images, we find that the leave-one-out accuracy was 92.1%, i.e. ca. 2,000 images were incorrectly labeled by the algorithm. We manually checked whether these were mislabeled or rather whether they “looked similar” in terms of area or landmark points in the photo, e.g. when both rosette or inflorescences were minute. We found that only 2.5% were incorrectly mislabeled (and corrected them) and are thus confident that the labeling error must be below 2.5%.

#### Experimental validation

Although repeating experiments in climatically-similar locations would be impractical, we could verify that survival in Madrid and low precipitation correlated with a preliminary drought experiment in the greenhouse^8^ (Spearman’s rho=0.17, n=211, *P*=0.01). On the other hand, reproductive allocation measured under optimal conditions in the greenhouse correlated with total seed output in the most similar field experiments, Tübingen high precipitation (Spearman’s rho=0.27, n=211, *P*=5×10^−5^)^66^.

Because we had two independent flooding tables per water treatment at each site, we also tested in an ANOVA the interaction field site and water treatment, the flooding table effect within location, and genotype effects. The results pointed to a replicability of the water factor, regardless of the two flooding tables, as well as its dominance in generating a fitness response in comparison to the effect of site (Table SII.2-3). These ANOVA tables were tested against the residuals. An even more conservative test for water vs site effects would be to treat the 16 spatial blocks (Fig. SII.1) as the unit of replication (rather than the plant pot replicate level). We conducted the F test of water, site, and interaction as F_water_ = MeanSq_water_ /MeanSq_block_ (and the same for site and the interaction). The results are most significant for the water treatment (*F*=918.4, *df1*=1, *df2*=12, *P*=1.0 × 10^−12^), then the interaction (*F*=110.3, *df1*=1, *df2*=12, *P*=2.1 × 10^−7^), and finally the site (*F*= 84.3, *df1*=1, *df2*=12, *P*=8.9 × 10^−7^). Note we use as degrees of freedom the corresponding 16 blocks minus the different treatments fitted. This confirmed once more the dominance of watering treatment over site treatment in determining plant fitness, although both as well as the interaction were important.

### VI. Author contributions

MEA conceived and designed the project. MEA carried out the experiment in Tübingen. MEA and RGR carried out the experiment in Madrid. All authors contributed to specific tasks in the experiments (see detailed description below). OB provided the field site in Tübingen and FGA provided the site in Madrid. DW secured funding for the project. MEA carried out the analyses and wrote the first draft of the manuscript. All authors edited, commented and approved the manuscript.

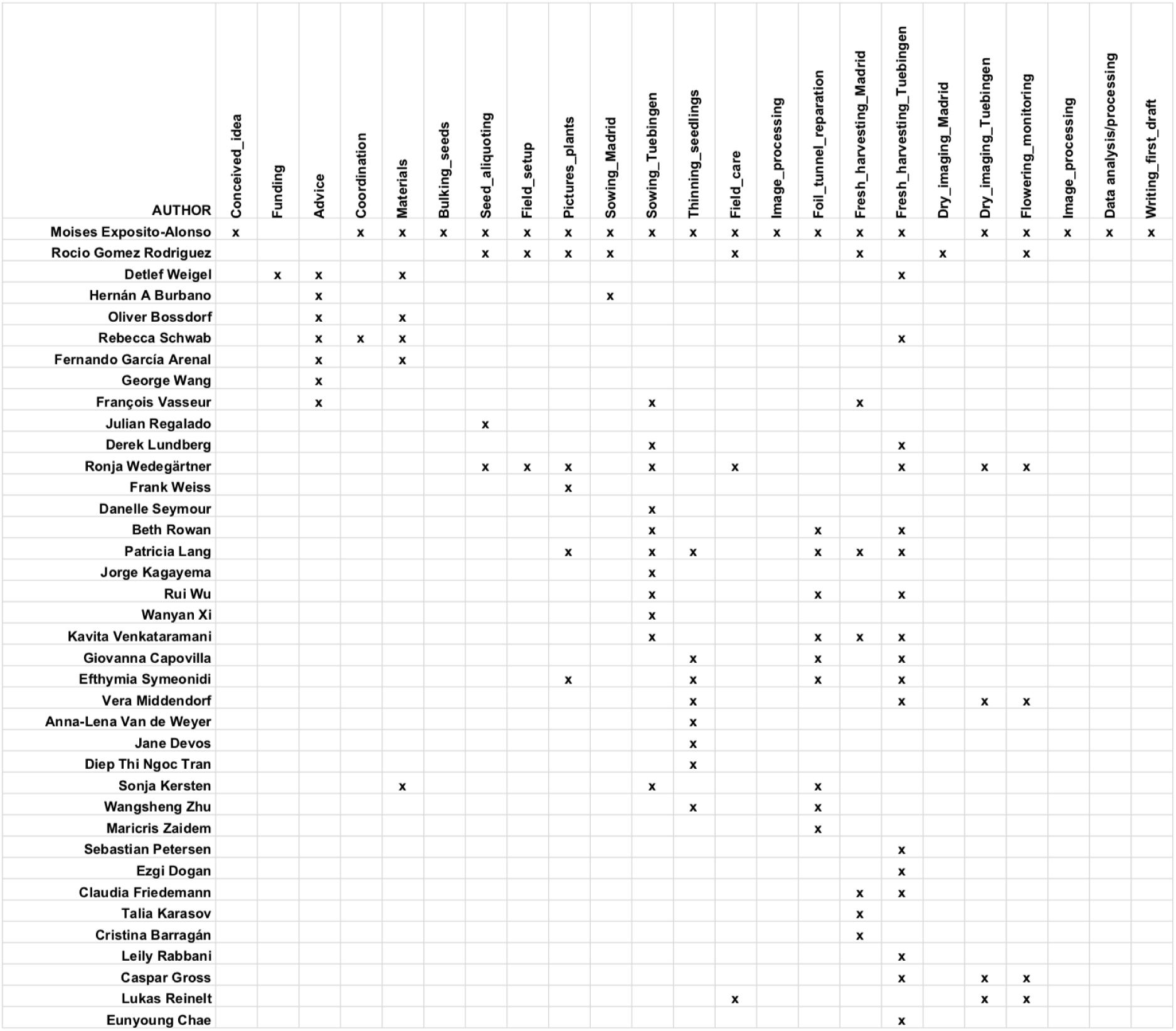

**Figure SII.1.**
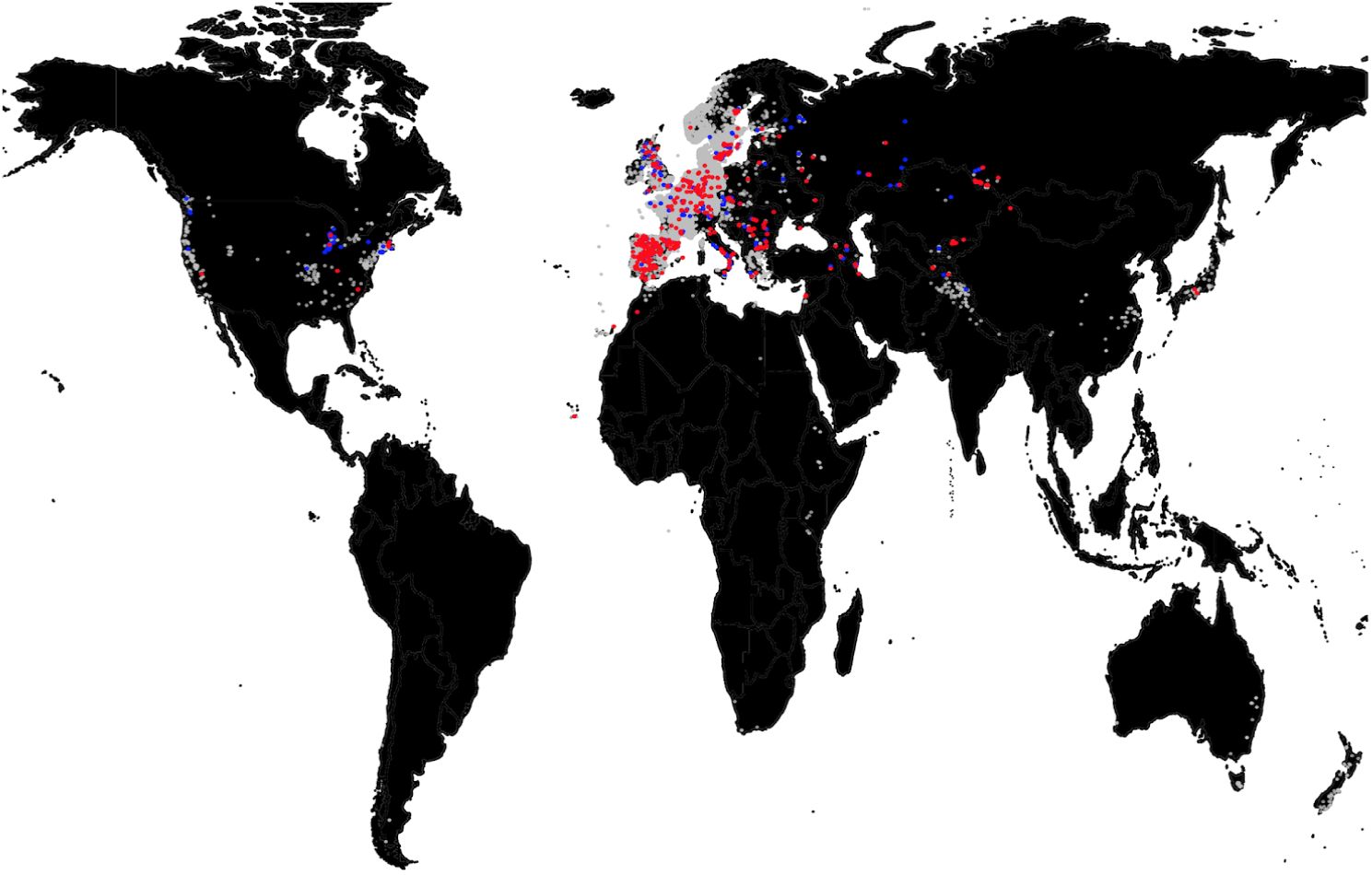
Geographic distribution of accessions. Locations of *Arabidopsis thaliana* accessions used in this experiment (red), 1001G accessions (blue), and all sightings of the species in gbif.org (grey).

**Figure SII.2.**
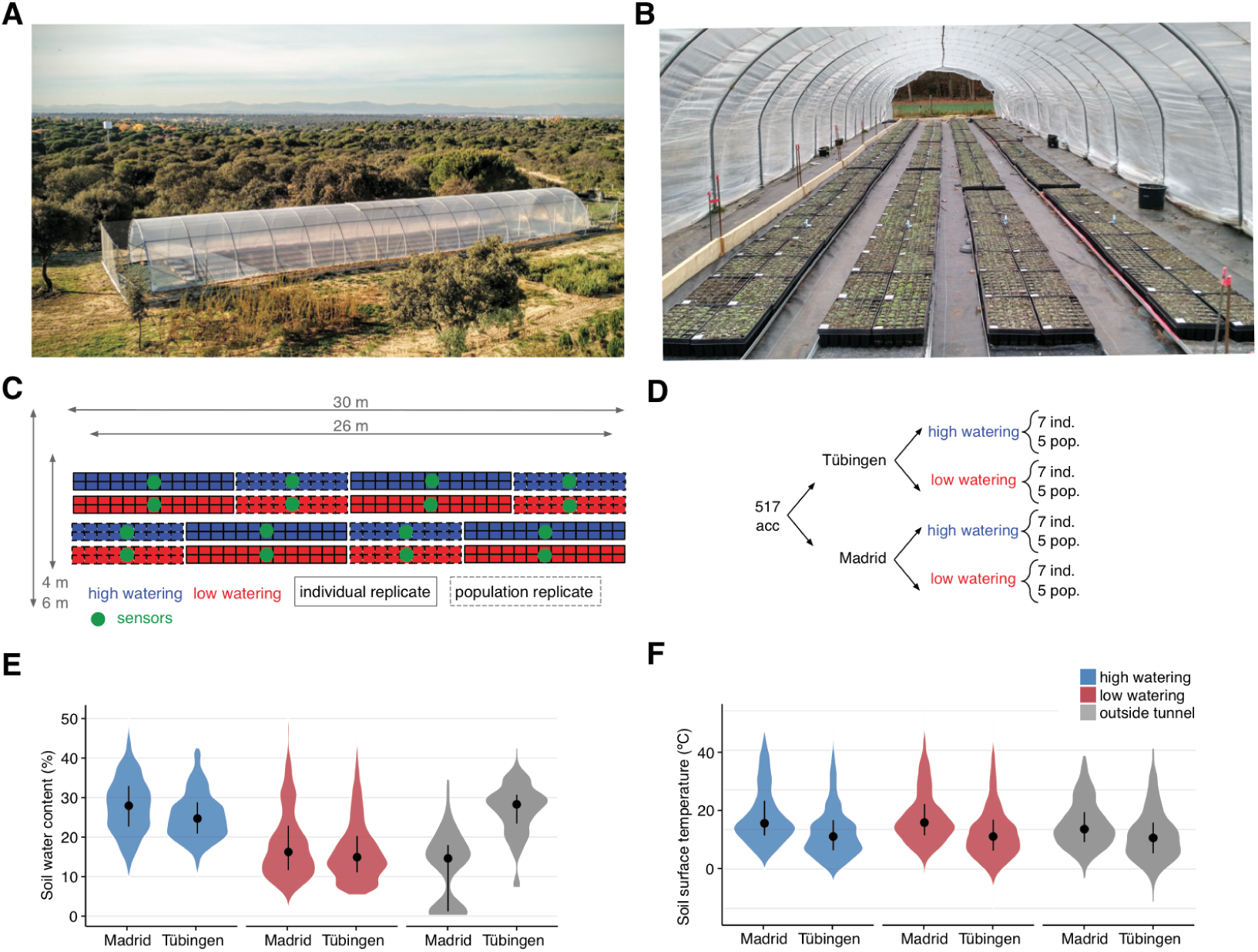
Field experiment design. (A) Aerial view of foil tunnel settings in Madrid and (B) view inside the foil tunnel in Tübingen. (C) Spatial distribution of blocks and replicates and (D) experimental design. (E) Soil water content and (F) soil surface temperature from the 34 sensors monitoring each experimental block and conditions outside the tunnel.

**Figure SII.3.**
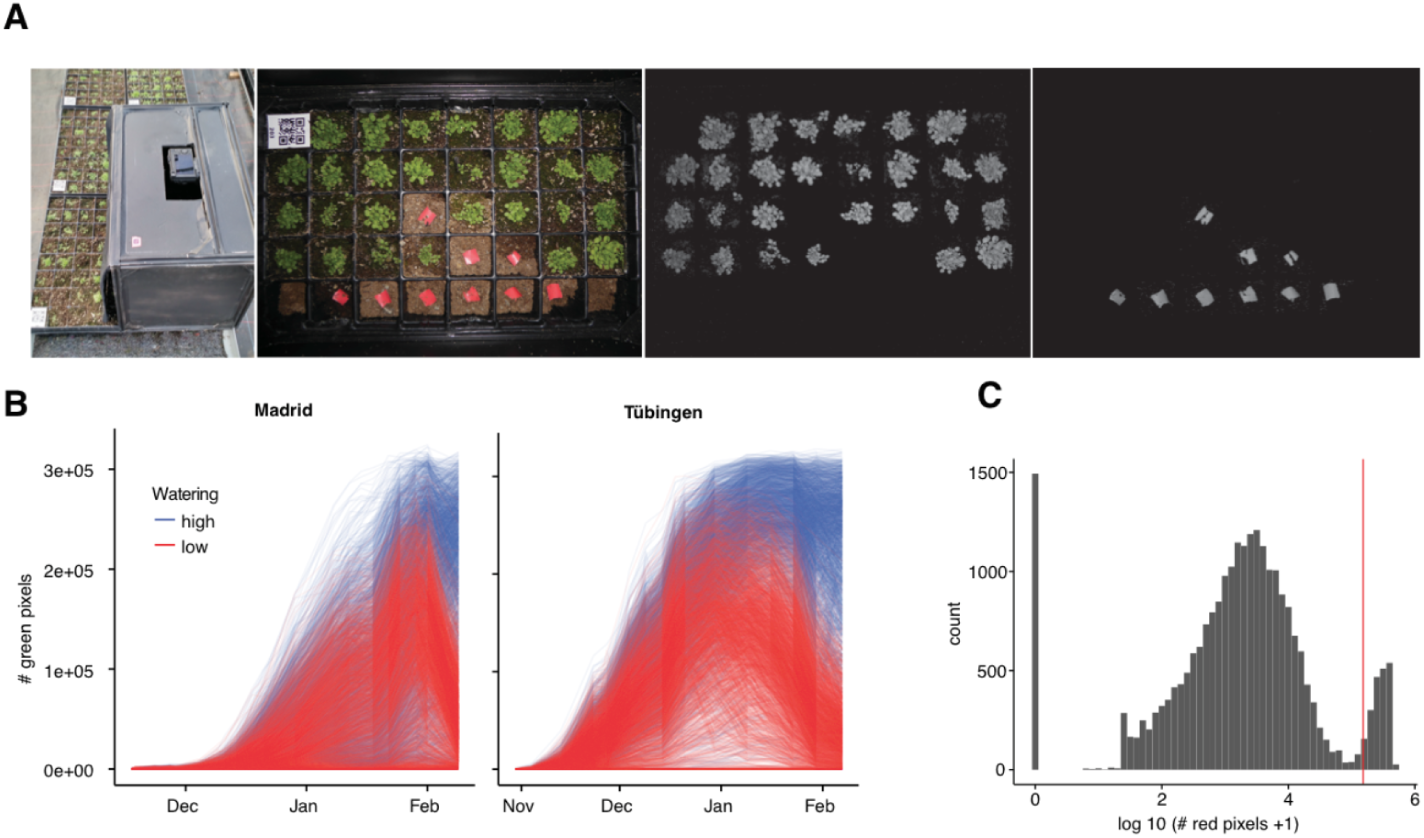
Rosette monitoring. (A) Customized dark box (“Fotomatón”) for image acquisition and example tray with the corresponding green and red segmentation. (B) Trajectories of number of green pixels per pot, indicating rosette area, for Madrid and Tübingen. (C) Distribution of the sum of red pixels per pot over all time frames. The red vertical line indicates the heuristically chosen threshold to define whether the pot actually had a red marker.

**Figure SII.4.**
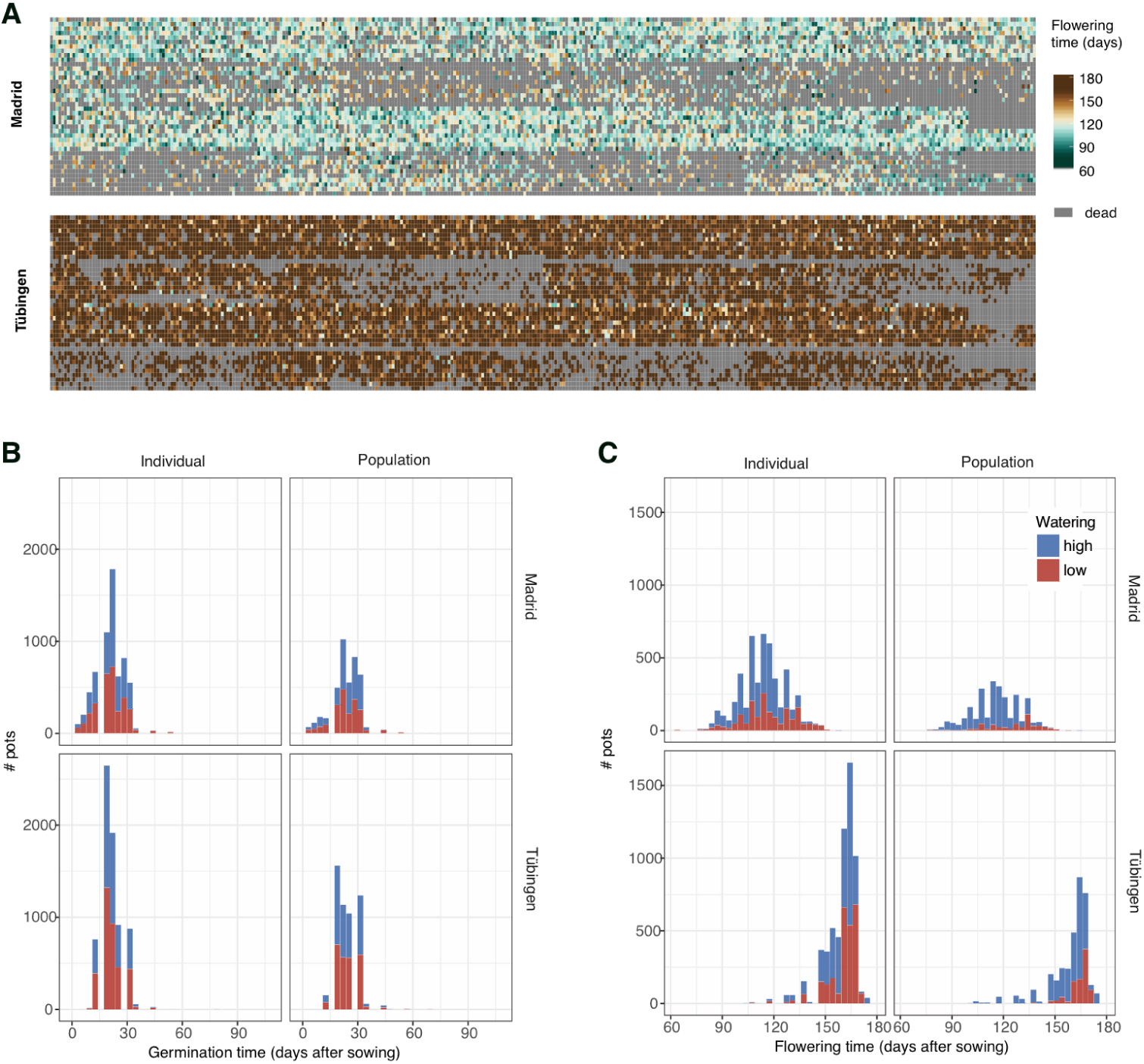
Flowering time distributions. (A) Flowering times per pot in the same spatial arrangement as in each tunnel (see Fig. SII.2). (B) Distribution of germination times. (C) Distribution of flowering times.

**Figure SII.5.**
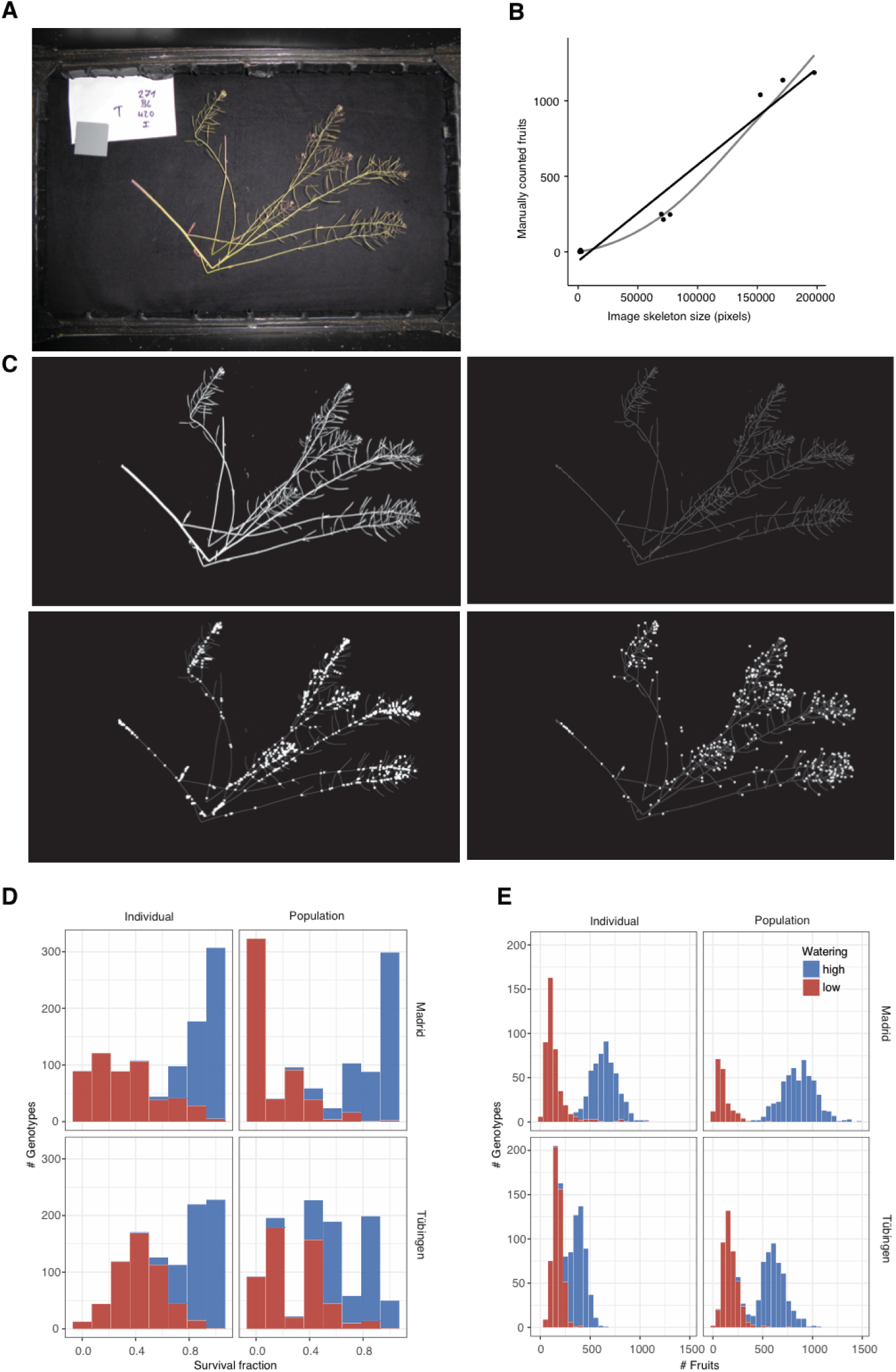
Inflorescence and seed set estimation. (A) Representative inflorescence picture. (B) Regression between the fruits of a few manually counted inflorescences and the inflorescence size calculated based on image processing. The four variables inferred in (C) accurately predicted the visually counted inflorescences as example (R^2^=0.97, n=11, P=10^−4^). (C) Resulting variables from image processing of (A): total segmented area (upper-left), skeletonised inflorescence (upper-right), branching points (lower-left), and endpoints (lower-right). Distribution of survival to reproduction (D) and fruits per plant (E) in the four environments.

---

**Table SII.1.** Summaries of environmental sensor measurements. A total of 34 sensors were placed in the different treatment blocks (low/high) as well as outside (out) of the foil tunnels. The median (interquartile) values of all sensors per treatment and location are shown.

| Site | Rainfall | Soil water content (%) | Air temperature (°C) |
| --- | --- | --- | --- |
| Madrid | out | 14.5 (1.09, 17.46) | 8.5 (5.34, 12.39) |
| Madrid | low | 16.1 (11.38, 22.51) | 10.0 (6.95, 15.13) |
| Tuebingen | low | 14.7 (10.76, 20.09) | 6.6 (3.27, 10.78) |
| Tuebingen | out | 27.7 (22.82, 30.50) | 5.6 (2.44, 9.54) |
| Tuebingen | high | 24.6 (20.73, 29.02) | 6.6 (3.27, 10.78) |
| Madrid | high | 27.8 (22.62, 33.00) | 9.8 (6.82, 15.13) |

**Table SII.2.**
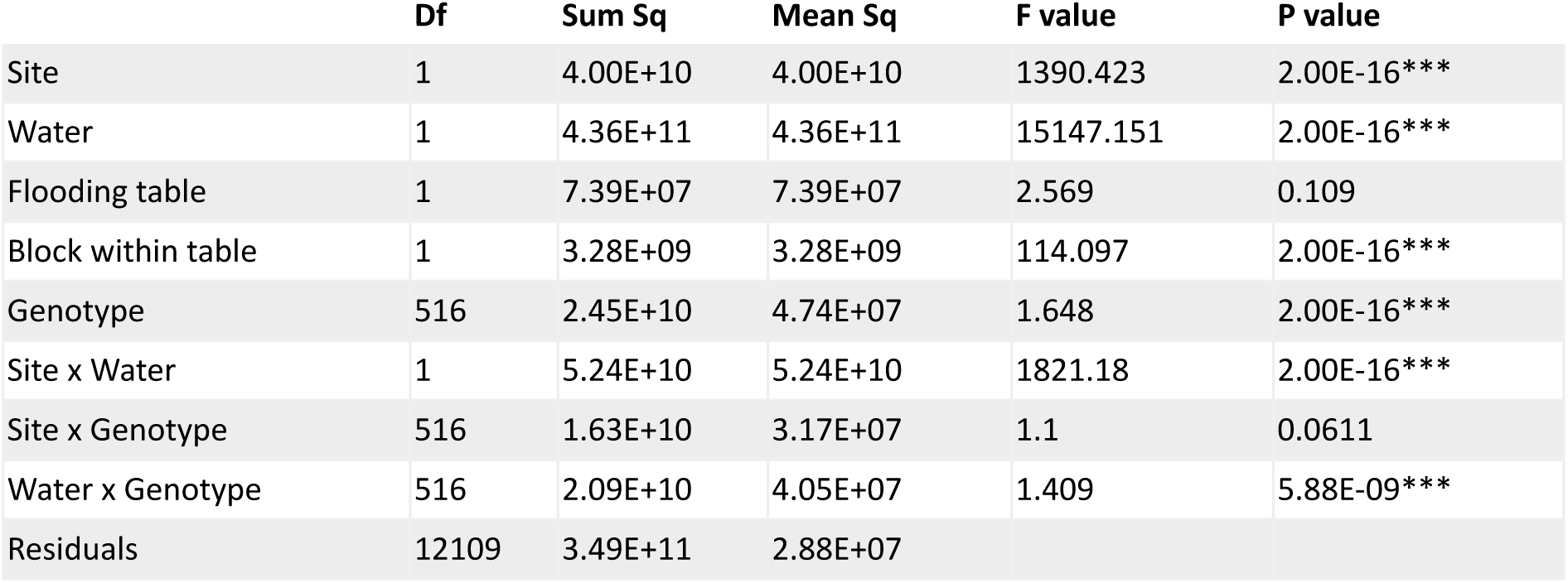
ANOVA table of fitness per individual. For the individual replicates, we run an AOV model in R of the expression: Fitness ∼ site + water + water:site + table + plot + genotype + genotype:site + genotype:water

**Table SII.3.** ANOVA table of fitness per population. Same as Table SII.2 but for population replicates.

|  | Df | Sum Sq | Mean Sq | F value | P value |
| --- | --- | --- | --- | --- | --- |
| Site | 1 | 56279467 | 56279467 | 983.256 | 2.00E-16*** |
| Water | 1 | 602839287 | 602839287 | 10532.179 | 2.00E-16*** |
| Flooding table | 1 | 14609050 | 14609050 | 255.234 | 2.00E-16*** |
| Block within table | 1 | 8794501 | 8794501 | 153.648 | 2.00E-16*** |
| Genotype | 516 | 40622483 | 78726 | 1.375 | 9.00E-08*** |
| Site x Water | 1 | 68696277 | 68696277 | 1200.19 | 2.00E-16*** |
| Site x Genotype | 516 | 32004305 | 62024 | 1.084 | 0.09969*** |
| Water x Genotype | 516 | 35219842 | 68256 | 1.192 | 0.00232 |
| Residuals | 7937 | 454296807 | 57238 |  |  |

**Table SII.4.**
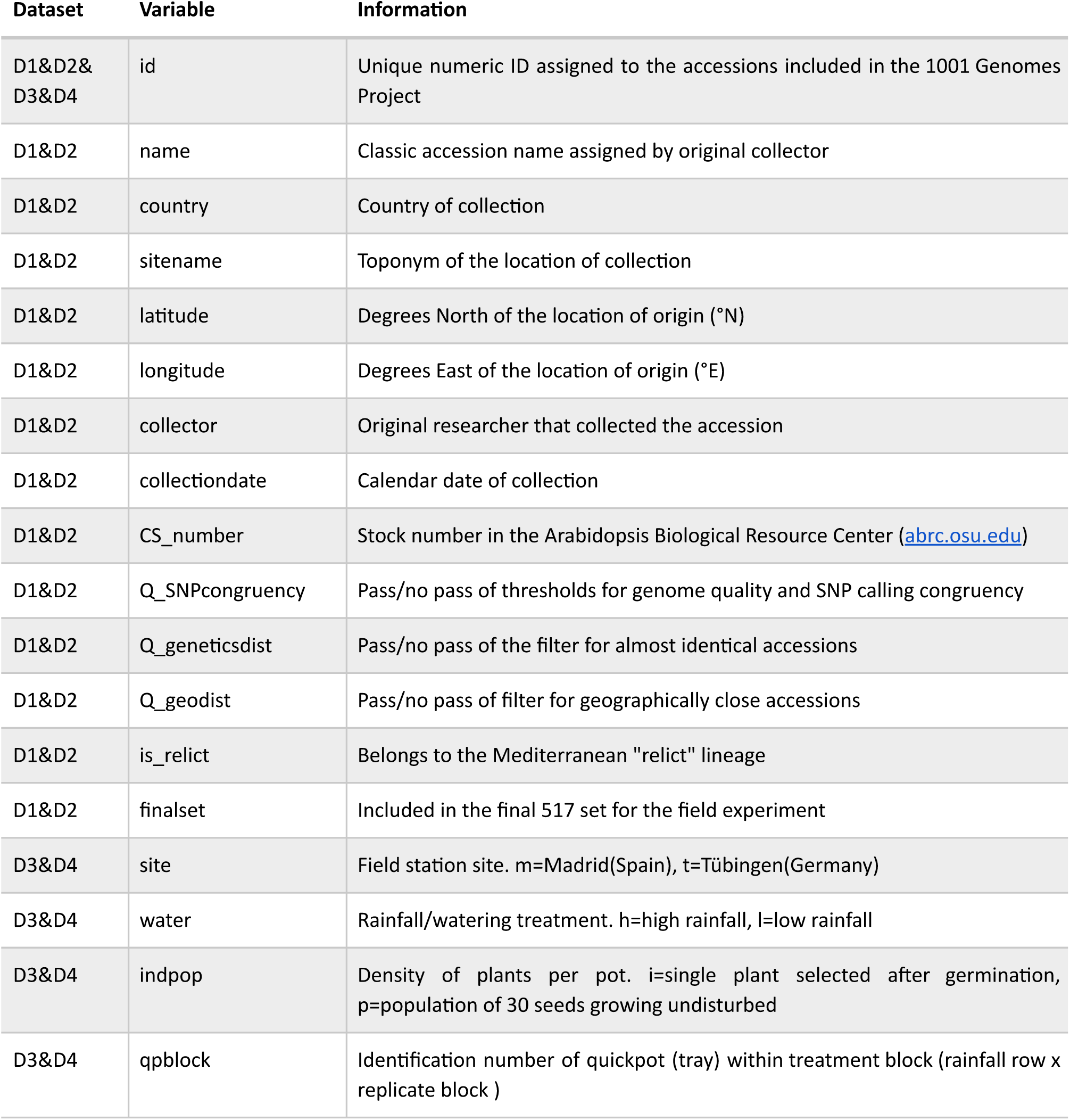
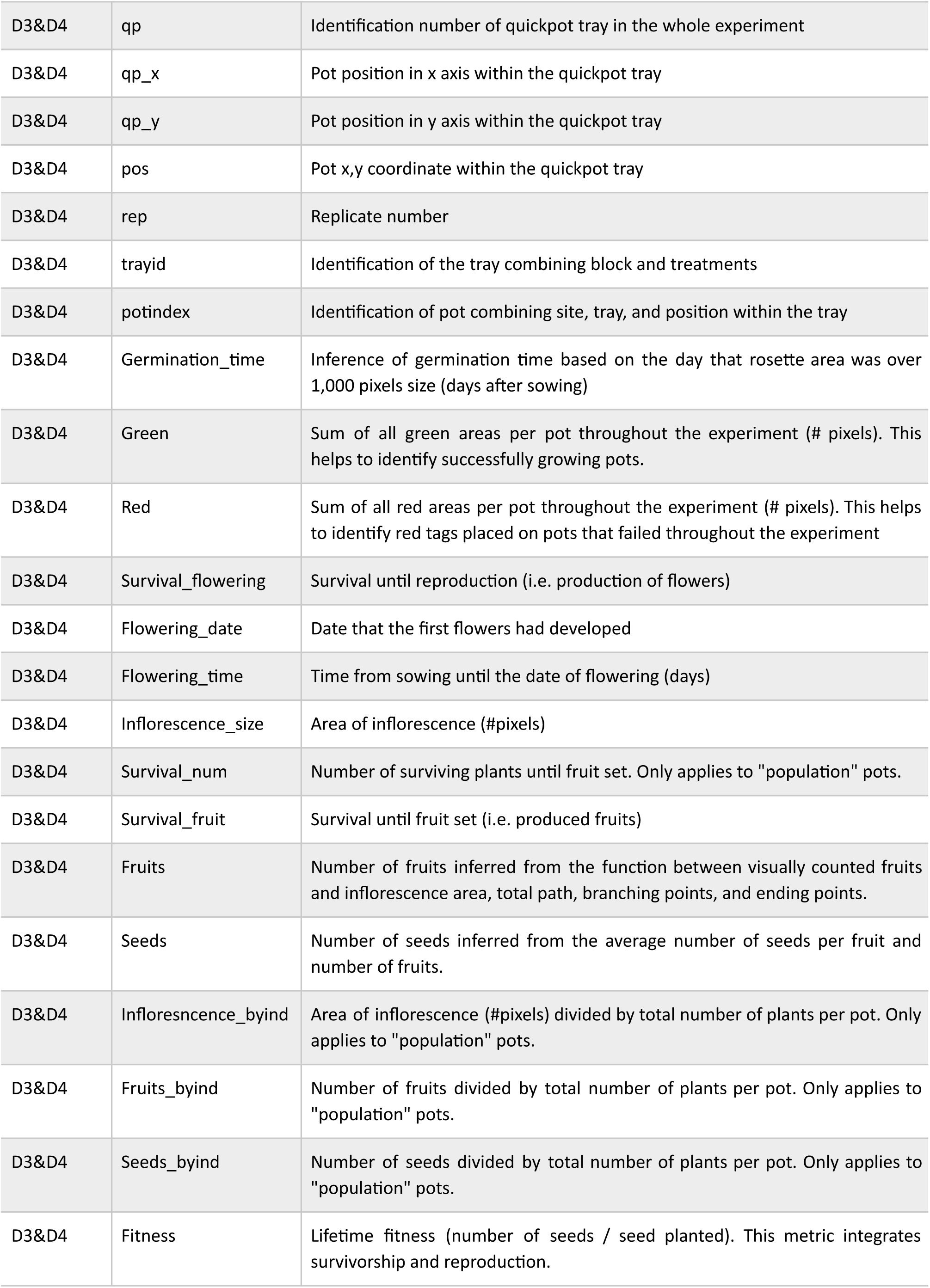
Variable descriptions. Variable names and their descriptions and units are reported (see <u>Datasets</u>). All datasets share a common accession identification number.

## DATASETS

Supplemental datasets are available in the online version of the paper [update for publication] and are also deposited at Figshare with doi: https://doi.org/10.6084/m9.figshare.6480599. A detailed description of each Dataset’s columns can be found in Table SII.4.

**Dataset 1 Quality-based selection of the original 1,135 accessions**

We report the 1001 Genome identification numbers, the quality filters that each accession passed during the selection of the 517 set.

**Dataset 2 Description of the 517 accessions**

We report the final set of 517 accessions that were used in the field experiment.

**Dataset 3 All traits measured per replicate**

For each pot replicate, we report all raw data as well as composite variables.

**Dataset 4 Curated means per accession**

For each accession, we report averages of all data as well as composite variables.

